# A neuro-computational approximation of the qualities of mental images

**DOI:** 10.64898/2026.08.25.747062

**Authors:** Rico Stecher, Daniel Kaiser

## Abstract

Mental images are challenging to study, given that our conscious experience is notoriously hard to access. The currently prevalent introspective methods are inherently subjective and can thus only provide limited access to their qualities. Here, we developed a neuro-computational approach that approximates and assesses the properties of mental images without the need for introspection. To enable this approach, we collected a large-scale EEG dataset (10 participants, 10 sessions each, 43,200 trials total) of participants imagining 16 scenes based on text prompts. We employed AI image generation to create candidate image sets that approximate the content of mental images (based on the imagined text prompts), computationally simulated visual cortex responses to these images and then assessed their representational alignment with rhythmic EEG responses during imagery. In line with previous reports, mid- to high-level features of the AI-generated candidate images yielded reliable alignment with human alpha activity. By manipulating the qualities of the candidate images, we then tested which qualities predisposed higher representational alignment with cortical imagery representations. We found an increased representational alignment for spatially blurred and low contrast images, providing evidence for the prevalent notion of a reduced sensory quality of mental images. We further found that mental imagery may be characterized by a psychedelic image style, which envelops the images in visual flows that distort the image proportions. These results show that our approach can objectively capture qualities of mental images without the need of introspection, providing a hypothesis-based alternative to emerging reconstruction approaches.

## Introduction

Mental imagery equips our brain with the ability to harness the sensory memory encoded during perception to simulate a seemingly endless set of internal experiences. A desire to grasp the elusive nature of our mental images has sparked more than a century of self-report studies^[1]^. Through these, we gained the prevalent view that mental images are typically described as of reduced sensory quality compared to actual vision and that they are characterized by differing degrees of vividness^[2,3]^.

In the prevailing view, our brain conjures mental images by reactivating visual cortex areas and representations that are also engaged during perception through top-down connections^[4]^. Recent research highlights that this top-down reactivation of visual representations is mediated by cortical alpha rhythms that carry information about imagined visual contents^[5–7]^. The representational geometry of these reactivated rhythmic visual representations can also be modelled with feedforward convolutional neural networks (CNNs; ^[5]^) trained for image recognition. We can thus gain insight into the visual features of mental images encoded in these rhythms using the same computational models we employ to characterize perceptual processing in the brain.

Despite these advances in understanding, a full picture of the qualities of our mental images is still elusive. While the reconstruction of perceived visual contents from neural recordings via generative approaches has recently made rapid progress^[8]^, directly reconstructing mental images is still challenging^[9,10]^. The most critical challenge is that imagery-related brain signals are inherently weak, likely because imagery only engages cortical layers that are involved in top-down processing^[11]^, which in turn results in a low signal-to-noise ratio (SNR) in the neural recordings of these signals. Such SNR limitations limit our ability to fully reconstruct mental images. In addition, previous reconstruction attempts have mainly focused on reconstructing the contents and not the qualities of mental images^[10]^. Yet, an objective estimate of mental images and their qualities would enable us to illuminate hard-to-verbalize characteristics of our conscious experience during imagery without relying on the currently predominant introspective reports that are highly subjective, prone to response biases, and have limited construct validity^[12–14]^.

Here, we introduce an approach that computationally approximates the contents of mental images and then assesses their qualities by evaluating how different image properties impact the alignment between AI-generated image features and imagery-related representations in the human brain. Our approach achieves this without the need for direct image reconstruction from neural activity. In our study, participants imagined 16 scenes based on short text prompts while we recorded their brain activity using EEG. To measure the weak imagery-related signals with sufficient statistical power, we collected a massive dataset of 10 participants performing 10 recording sessions each, amassing a total of 43,200 trials. We computationally approximated the contents of participants’ mental images by feeding each imagined prompt into a latent text-to-image diffusion model (LDM; ^[15]^), an AI algorithm that can generate images based on text input, to generate large sets of candidate images for each prompt. The resulting image sets were then evaluated by a feedforward CNN trained for scene recognition. Finally, we employed representational similarity analysis (RSA; ^[16]^) to determine if the representational geometry in the evaluator CNN matched the representational geometry in EEG alpha-band activity recorded from the human participants. We found a representational correspondence between participants’ alpha-band representations in intermediate and late processing stages of the CNN, suggesting that our generated image sets were indeed sufficient as an approximation of the contents of their mental images.

Having established that our generated image sets suffice as an approximation of our participants’ mental imagery contents, we sought to objectively test their qualities. We manipulated the images’ properties to test whether creating image sets that are more akin to participants’ imagery enhances the representational alignment between the evaluator CNN and participants’ alpha-band activity. To create image sets that approximate the qualities of mental images, we either altered their image features in targeted ways or used an LDM fine-tuned for art style transfer to alter their style.

In the image feature manipulation, we found that degrading the images by lowering their contrast or spatially blurring them increased the representational alignment. This provides objective evidence for the widespread intuitive notion of visual imagery being an impoverished sensory experience. In the style manipulation, we observed tentative evidence that mental images may be characterized by a psychedelic style which envelops the images in shapeless visual flows that distort the image proportions, suggesting that mental imagery may be characterized by such flows. Overall, our approach demonstrates the feasibility of capturing the qualities of mental images without the need for subjective introspection.

## Results

We recorded the EEG activity of 10 participants (6 female) imagining 16 scenes according to short prompts for 10 sessions each (43,200 trials in total). Prompts were presented for 3,000 ms and participants imagined the scenes for 4,000 ms.

### AI-generated image sets approximate the contents of mental images encoded in large-scale human brain data

We approximated the imagined visual contents of the participants by feeding each of the 16 imagined scene prompts into Stable Diffusion 2.0 (SD; https://huggingface.co/docs/diffusers/en/api/pipelines/stable_diffusion/stable_diffusion_2), a LDM^[15]^ which generates images based on text. For each prompt, we generated 100 candidate images (i.e., 1,600 images in total). We hypothesized that if we can generate a distribution of images large and diverse enough, the participants’ actual imagined visual contents should fall within this distribution.

To assess the alignment of the image set with the visual contents represented in the neural activity of the participants, we first simulated visual cortex responses to this image set using VGG-16^[17]^, a CNN model, trained for scene recognition using Places365^[18]^. To probe the representational geometry of the CNN, we extracted its unit response patterns at each layer and created representational dissimilarity matrices (RDMs; ^[16]^) by computing a correlation distance between the response patterns of each pairwise combination of the 16 images. This was done separately for each of the 100 image versions, and RDMs were then averaged across image versions. Since there is an inherent autocorrelation in neighboring CNN layer RDMs, we averaged layer RDMs into 4 layer group RDMs (early, early intermediate, late intermediate, fully connected; ^[5]^), yielding a quantification of the representational geometry at 4 stages of increasingly complex visual feature coding.

To compare the representational geometry in the CNN to the representational geometry in the human brain, we next quantified the similarity in EEG alpha power patterns during imagery for our human participants. We first reduced session-by-session variation in the EEG data by mapping the power pattern space to a feature space that both is maximally independent from all sessions and maximizes the discriminability among scenes via semi-supervised maximum independence domain adaptation (sMIDA; ^[19]^). We then constructed an alpha band RDM from pairwise correlation distances of the domain-adapted alpha activity patterns, using the frequency with the greatest mean pairwise distances in the 8-13 Hz range. Finally, we quantified the representational alignment between the evaluator CNN and the human alpha band by correlating the CNN layer group RDMs with each participant’s alpha band RDM. The correlations at each layer group were tested against 0 with a right-tailed one-sample t-test (N = 10, p < 0.05) and the resulting p-values were FDR-corrected across layer groups.

In our initial analysis, we investigated which level of image variability is optimal for approximating the content of mental images by generating 3 different image sets with decreasing LDM prompt adherence, which we achieved by systematically lowering the classifier-free guidance scale model parameter (cfg; ^[20]^), which steers this property. We chose the cfg values 7 (medium variability), 3 (high variability), and 1 (very high variability), since they resulted in image sets with distinct variability levels (see Figure 2A) and conducted this analysis separately for each set (see Figure 2B).

We found positive correlations that were significantly greater than 0 from the intermediate to fully connected layer groups for all cfg values. For cfg 1, these correlations ranged from the early intermediate to the fully connected layer group (peak p = 0.007) and for cfg 3 (peak p = 0.006) and 7 (peak p = 0.02), from the late intermediate and the fully connected layer groups. There were no meaningful correlations in early layers across the image sets and these were also the most variable across participants. This suggests that the generated images were a sufficient approximation of the participants’ imagined visual contents and that alpha rhythms encode mid- to high-level visual features of the imagined scenes^[5]^.

Looking across the different cfg values, we found a trend towards greater representational alignment with greater image variability. The general correlation pattern was, however, qualitatively similar across cfg values. As each image set was generated with new random seeds, these results further show that our findings generalize across distinct image sets.

### Targeted manipulation of image properties reveals the qualities of mental images

Having established that the generated image sets suffice as an approximation of our participants’ mental imagery contents, we next sought to manipulate key characteristics of the generated images and test whether these manipulations could enhance the alignment with human mental imagery. In all following analyses, we altered the properties of the image set for which we observed the most pronounced representational alignment with the participants’ neural imagery data (cfg 1). We conducted an image feature manipulation and a style manipulation.

The aim of the image feature manipulation was to capture the impoverished quality typically assigned to mental images^[1,3]^. Based on our participants’ Vividness of Visual Imagery Questionnaire (VVIQ; ^[2]^) scores, their imagery is of average vividness with no signs of hyperphantasia (scale-reversed VVIQ mean = 55, range: 48-61) suggesting that their mental images should be of this impoverished quality, too. We attempted to emulate this quality by creating 4 image sets to which we applied 1 of 4 types of image degradation (grayscale, blur, noise and low contrast respectively) and 1 vivid (high saturation, high contrast) control set which should be more dissimilar to the quality of their mental images (see for Figure 3A & B for examples).

For each altered image set, we had VGG-16 evaluate the images, created layer group RDMs as before, and predicted the alpha band RDMs of each participant with a regression model, using the layer groups for which we found a correspondence with the neural imagery data in our previous analysis (early intermediate, late intermediate and fully connected) as predictors. We then compared the average explained variance (R²) across participants of each altered image set to the average explained variance of the original cfg 1 image set as a baseline condition. An increase in R² would suggest that the property manipulation rendered the properties of this image set more alike the properties of our participants’ mental images. This was tested for significance with a right-tailed paired Wilcoxon signed rank test (N = 10, p < 0.05) and the resulting p-values were FDR-corrected across comparisons.

For the image feature manipulation, we observed a statistically significant increase in R² relative to baseline for the blurry (p = 0.034) and low contrast (p = 0.034) image sets. We found no significant effects for any of the other conditions. For the vivid image set, we saw a minor drop in R² compared to baseline (did not reach significance in an exploratory two-tailed test), which supports the notion of it being more dissimilar to our participants’ mental images, at least on the descriptive level. These results suggest that reducing the image quality by blurring the images or reducing their contrast results in a better fit with the imagery data. Taken together, our data showcases that our approach cannot just approximate the contents of visual imagery, but can capture its reduced sensory quality.

The aim of the style manipulation was to alter the style of the image set to art styles that may have inherent properties that are akin to some properties of mental images (see Figure 3A & C for examples). The first style we investigated was a psychedelic style, since it envelops the image set in shapeless visual flows that distort the image proportions, similar to what is commonly experienced during hallucinations^[21]^. We were interested if our participants’ mental images are characterized by such flows. The second style we examined was a watercolor style, since watercolor images tend to be faint, with blurred and undefined outlines, which have previously been described as phenomenological qualities of mental images^[1]^ . The third style we assessed was a surrealist style. A lot of surrealist art tends to be characterized by abstract, unrealistic and disproportionate scene arrangements. We wanted examine if this style can capture the bizarre aspects of dreaming and waking imagery that people have reported to experience^[22]^. For the purpose of positive control, we created some image sets in which we altered the style, but chose styles that we believe do not have any imagery-like properties. We decided on the following styles: 3D model, cubism, pixel art. The style transfer was carried out with DreamShaper 8 (https://civitai.com/models/4384/dreamshaper), an SD-1.5-based model fine-tuned for generating art styles.

For the style manipulation, we found a significant increase in R² for the psychedelic style (p = 0.041). We did not observe any significant increases for any of the other styles. All observed relative decreases in R² were not significant in an exploratory two-tailed test. These results suggest that imagery may be characterized by shapeless visual flows that distort the image proportions. However, when reconducting the style analysis with a VGG-16 architecture trained on stylized object images (Stylized-ImageNet; ^[23]^, see supplementary Figure S3) to investigate if the lack of representational alignment for the other styles may be due to the images being out of distribution of the evaluator CNN, we were unable to find any effects for the styles. This suggests that the psychedelic effect might not be as robust and that the absence of an effect for the other styles could not be explained by the images being out of distribution for the evaluator CNN.

## Discussion

Here, we developed an approach that approximates mental images by leveraging AI image generation to create large image sets of candidate images and a CNN to simulate visual cortex responses to these images. By evaluating the representational alignment between this CNN and visual content representations encoded in cortical alpha rhythms during mental imagery in humans, we could benchmark which properties of an image set predispose greater alignment and thereby infer the qualities of mental images.

We evaluated this approach by collecting a massive visual imagery EEG dataset (10 participants, 10 sessions each, 43,200 trials in total). We found that the generated image sets yield an effective approximation of the contents of participants’ mental images. Furthermore, we show that targeted manipulations of the images’ properties can be harnessed to objectively test the qualities of mental images.

When comparing representations for the generated image sets (using a CNN as an evaluator) to representations of imagined scenes (using alpha activity in the EEG), we found a representational correspondence between participants’ alpha band imagery representations in intermediate-to-late CNN processing stages. This effect was found across different image sets with different variability constraints, suggesting that the generated image sets are a reasonably reliable and practicable approximation of participants’ imagined visual contents.

### Quantifying the feature complexity of mental images

The reliable correlations in intermediate and late processing stages of the CNN suggest that reactivated visual features encoded in alpha rhythms carry mid-level and complex high-level features of imagined scenes. This result is consistent with findings from object imagery, where alpha band representations during imagery are predicted by high-level features extracted by a CNN trained for object recognition^[5]^. It is further consistent with alpha rhythms carrying mid- to high-level scene information during imagery, such as whether the scene is natural or man-made, cluttered or sparse, and open or closed^[7]^. In this study, we showcased that, by AI-generating the imagined visual contents and computationally simulating the perception task with a CNN, the feature complexity of rhythmic scene representations during imagery can be characterized without any visual reference in the imagery paradigm and without the need for a separate perception task in the experiment^[5]^.

### Quantifying the quality of mental images

The key promise of our proposed approach, however, lies in its ability to objectively quantify the qualities of mental images by manipulating image properties in a targeted way. To achieve this, we either applied image filters or prompted the generative network to create images of a certain quality and tested whether images with these qualities in turn yield a better approximation of neural representations during imagery. This approach goes beyond the state of the art in the field, where the qualities of mental images are inferred through introspective insights captured by descriptions^[24]^, single-item-rating scales^[25]^ and questionnaires (such as the VVIQ; ^[2]^). While a lot of the reported qualities do converge and have enabled us to roughly characterize how people would describe the qualities of their mental images^[1]^, it is still not clear how these descriptions map to their actual experience due to the inherent subjectivity and insufficient validation against objective indicators of the prevailing methods^[14]^. Our approach complements these methods by providing an objective neural benchmark of the qualities of mental images, which has the potential to validate introspective insights or identify novel hard-to-verbalize qualities.

Here, we altered the qualities of images in two targeted ways, manipulating image features and style. In the image feature manipulation, we found a higher representational correspondence compared to the more naturalistic baseline image set for the blurry and low contrast image sets. This finding aligns with previously discussed dimensions of imagery vividness. Blurriness/sharpness and contrast have been postulated as dimensions contributing to the vividness of mental images by multiple authors^[1,26,27]^. According to Fazekas^[26]^, blurriness and contrast are key dimensions that determine both the subjective intensity (i.e., vividness) and specificity of mental images. This means the specific degradations we found an effect for have previously been discussed as key aspects of the reduced sensory quality of mental images in the imagery vividness literature. Our approach seems to be able to capture such properties without the need for introspection. Surprisingly, we found no effect for the grayscale set, though previous literature has also discussed colorfulness (or lack thereof) as an important factor^[1]^.

For the style manipulation, we observed group-level evidence of mental images being characterized by the psychedelic style. Based on the results in the image feature manipulation and our participants’ only average reported vividness, we believe that this effect is not driven by the very vivid color scheme it induces, but rather by how it envelops the images in these shapeless flows that distort the image proportions. Imagery can fluctuate^[28]^ and its contents tend to constantly change in unpremeditated ways, which has previously been termed “metamorphosis”^[1]^. It may be that this effect is caused by fluctuations in participants’ mental imagery and how they distort the proportions of the mental image. However, due to how marginal this effect is and the fact that we could not replicate it with VGG-16 trained on Stylized-ImageNet (although this may also have been due to a generally reduced representational alignment of this model with the imagery data we observed, potentially because the pre-trained model was not properly optimized), we would only interpret it with caution. It does, however, highlight the potential of our approach in identifying unexpected properties of mental images.

We observed no group-level evidence of our participants’ mental images being characterized by any of the other art styles we tested. The most apparent explanation would be that our participants’ mental images were not characterized by any of the properties inherent to these styles. It’s also possible that the participants did experience some of the characteristics that we wanted to capture through the style manipulation, but the way the model implemented those characteristics didn’t match the participants’ specific visual experience.

In this study, we provided a foundation for objectively testing the properties of mental images via image manipulation. However, it is not feasible to exhaustively test all potential properties of mental images in a single study. Future studies are now open to broaden the range of properties we tested. Other properties that may be worth investigating are for example brightness^[1]^ and opacity^[13]^.

### Limitations

A first limitation is the relatively small sample we tested. We consciously made the trade-off to collect fewer participants, but gather as much data for each individual participant so that we could measure their representational geometry as accurately as possible. The resulting small sample size may, however, limit the generalizability of our conclusions to the general population. In addition, our participants’ imagery is of average vividness and doesn’t cover the whole vividness spectrum. While this shows that exceptional imagery abilities are not required for this approach to be effective, it would be enlightening to see how vividness at the highest and lowest percentiles of the spectrum would modulate our results. We thus deem the application of this approach to a larger sample with more diverse imagery abilities to be a fruitful direction.

Despite our best efforts at collecting as much data as possible to maximize the power for every participant, the SNR in our data was poorer than we expected. This is why we had to resort to 1) extensive data preprocessing, 2) averaging to improve the SNR and 3), group-level rather than individual-participant analyses. Whether the prevailing low SNR is related to concrete design or analysis choices in our study or whether it alternatively reflects hard limits in EEG data acquisition remains, at this point, unclear.

A further limitation of this study was the limited number of stimuli. We kept the stimuli at 16 scenes to maximize the number of trials per scene and thus maximize the SNR in the neural responses for each scene. Future studies could try employing this approach with a large and variable set of stimuli (also covering other visual domains) to demonstrate its ability to generalize across different imagined visual contents.

Due to the nature of our pipeline, we have no direct subjective evaluation of how similar our generated images are to our participants’ actual images. Our approach centers around capturing the distribution of imagined visual contents through creating large sets of candidate images (1,600 images for each of the 14 image sets we analyzed here) and then integrating these estimates on a very abstract and hard-to-visualize level (the representational geometry of the CNN’s unit responses to each set of image versions). This makes a direct similarity rating on the image level or on the integrated level by our participants somewhat infeasible. However, given that we found robust group-level effects, the generated image sets were sufficiently capturing participants’ mental images. In this context, it is important to note that we do not claim that the generated images are our participants’ actual mental images. Yet, they were close enough to our participants’ actual mental images to approximate imagery-related representations.

## Conclusion

In sum, we present a novel approach that can computationally approximate mental images and evaluate their alignment with human brain responses during imagery. This approach allows for a targeted investigation of the properties of mental images, providing a hypothesis-driven alternative to emerging reconstruction approaches. Our study provides an objective test of the properties of participants’ mental images, providing evidence of the impoverished nature of mental imagery. We further provide tentative evidence of mental imagery being characterized by shapeless visual flows that distort the image proportions. More generally, our approach establishes a starting point for a detailed mapping of the properties of mental images without relying on subjective reports.

## Methods

### Participants

12 participants took part in this study. One participant was excluded because their English proficiency was not sufficient. Another participant was excluded because they did not finish all 10 EEG sessions. In total, 10 participants (6 female, mean age = 24.7, SD = 5.1) with all 10 sessions completed were included in the analysis. Participants were compensated with 10€ per hour and received a 50€ bonus after finishing the 10th session.

We assessed participants’ imagery ability in advance using the Vividness of Visual Imagery Questionnaire (VVIQ; ^[2]^). Participants were only included if their VVIQ score exceeded a cut-off of 32 (scale reversed; as described in ^[29]^) to ensure that they did not suffer from aphantasia. All participants provided written consent. The study was approved by the Ethics Committee of the Justus Liebig University Giessen and was in accordance with the Declaration of Helsinki.

### EEG task and procedure

Every participant completed 10 EEG sessions. Each session consisted of the initial preparation of the EEG cap, followed by a computer experiment in which participants had to imagine natural scenes based on short prompts. In the first session, we included a practice run that preceded the main experiment. Here, participants went through 16 practice trials in which they imagined each of the scene prompts once to familiarize themselves with them. The scene imagery task (Figure 1) was an adjusted version of the task in our previous study^[7]^. On each trial, participants were initially presented a scene prompt at the center of the screen surrounded by a black frame (21° horizontal visual angle). The frame stayed on the screen for the entire trial. Participants had 3,000 ms to read the prompt, after which a black fixation dot appeared at the center of the screen. After a randomized interval between 1,000-2,000 ms, the fixation dot turned red, which served as the cue for the participants to imagine the scene within the frame while keeping their gaze fixated on the dot. The dot stayed red for 4,000 ms and participants were instructed to maintain their mental image for the entire period. The dot then went back to black. Trials were separated by an intertrial interval varying randomly between 800-1,200 ms. The main experiment consisted of 27 blocks of scene imagery trials. In each block, participants had to imagine each of the 16 scene prompts in random order. Participants were allowed to take frequent self-paced breaks. They thus completed 27 x 16 = 432 trials of the scene imagery task per session, resulting in a total of 10 x 432 = 4,320 trials per participant across all sessions (43,200 across all 10 participants). They also filled in an online questionnaire on LimeSurvey (www.limesurvey.org) at the end of each session in which they were asked to rate the properties (openness, naturalness, clutter level) of each imagined scene on a scale from 1-7 to ensure that they had imagined the scenes at least roughly as intended. These ratings were as expected. Each session was typically between 2.5-3.5 h long. The experiment was run using PsychToolbox^[30]^.

**Figure 1.**
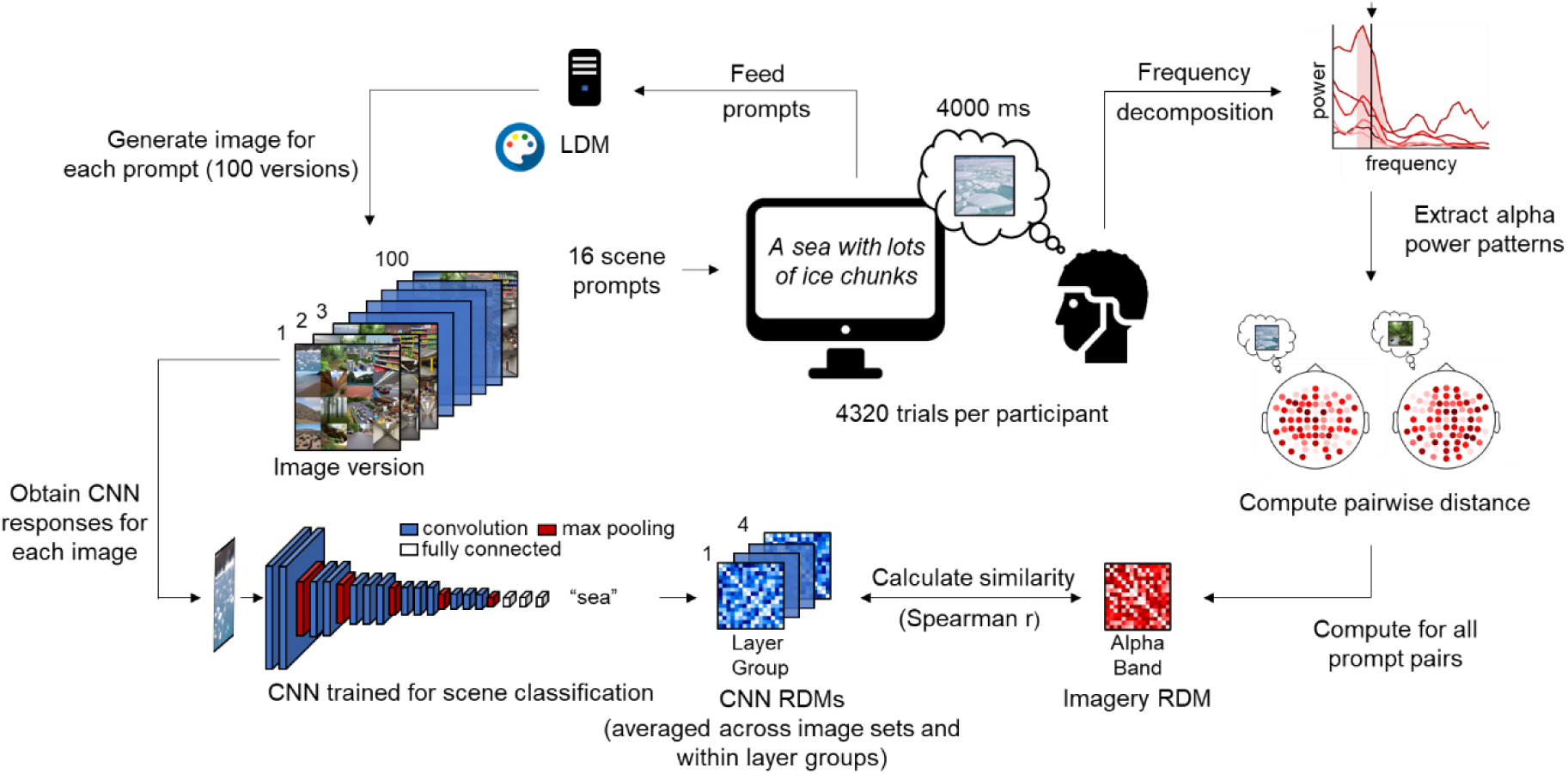
Pipeline for approximating imagined visual contents and evaluating their alignment with rhythmic neural representations of mental images. On each trial, participants imagined 1 of 16 scenes for 4,000 ms, based on short text prompts, while their EEG was recorded across 10 sessions (4,320 trials per participant). To approximate imagined visual contents (left hand side), we fed each of the 16 text prompts to a latent text-to-image diffusion model (LDM) and generated 100 different candidate images. This yielded 100 image version sets (i.e., 1,600 images in total). A convolutional neural network (CNN) trained for scene recognition then evaluated these image sets and we extracted representational dissimilarity matrices (RDMs) for 4 layer groups along the CNN hierarchy. To establish the neural similarity among mental images (right hand side), we quantified the representational geometry of the imagined visual contents in the EEG by creating RDMs from alpha band power patterns across electrodes. Finally, we correlated the CNN RDMs and the alpha band RDMs to establish their representational alignment.

### Stimuli

Participants were presented 16 short prompts describing natural scenes (3-7 words; see supplementary Figure S1 for an overview of all prompts). To cover a range of relevant perceptual properties, the scenes were chosen to vary in their openness (8 open and 8 closed scenes), naturalness (8 natural and 8 man-made scenes) and clutter level (8 cluttered and 8 sparse scenes).

### EEG preprocessing

EEG data was acquired using an Easycap system with 64 channels and a Brain Products amplifier. The data was recorded at a sample rate of 500 Hz with Fz as the reference. The electrode arrangement followed the standard 10–10 system. All preprocessing was conducted using FieldTrip^[31]^. The EEG was band-stop filtered to remove 50 Hz line noise, epoched between -1,000 ms and 5,000 ms and baseline-corrected with a baseline window of 500 ms. The EEG signal was then downsampled to 200 Hz. Noisy channels were removed by calculating the variance of each channel and rejecting outlier channels on this metric through visual inspection. Finally, independent component analysis (ICA) was applied to the EEG data and eye artifact components were removed through visual inspection.

Missing channels were interpolated by calculating the average of the surrounding channels within each EEG session in order to keep the same number of features across sessions.

### Frequency decomposition

The potential at each EEG electrode during the entire 4,000 ms imagery period was decomposed into 6 discrete frequencies ranging from 8-13 Hz (1 Hz frequency resolution) for every trial. We used multitapers (15 dipolar spherical sequence tapers) with 2 Hz frequency smoothing to boost the SNR in the inherently noisy imagery EEG data^[7,32]^. The frequency decomposition was conducted using FieldTrip.

### Analysis pipeline

#### Image generation

We leveraged AI image generation to approximate the participants’ imagined visual contents by generating a diverse distribution of possible imagined scene images. The images were generated by feeding each of the imagined scene prompts into a LDM^[15,33]^. Specifically, we used Stable Diffusion^[15]^ 2.0 implemented in the Automatic1111 WebUI (v.1.6.0; https://github.com/AUTOMATIC1111/stable-diffusion-webui) running in Python 3.10.6. This model operates as follows.

During forward diffusion, noise is added to images step-by-step and a neural network (a UNet) learns to predict how much noise was added at a particular step (relative to the original image). Images are then generated by reversing this process (reverse diffusion) and using the neural network to denoise a noise image step-by-step. Aligning the denoised image with the supplied text input is achieved by using a CLIP^[34]^ text encoder to obtain text embeddings and conditioning the neural network on these embeddings via its cross-attention layers. This diffusion process is made more computationally efficient by first using the encoder of a variational autoencoder to encode the high-dimensional pixel space of the images into a more low-dimensional latent space and conducting the diffusion on this latent space. Once this is complete, the decoder of the variational autoencoder then converts the low-dimensional space back to the high-dimensional pixel space.

Images were generated on the CUDA cores of an NVIDIA® GeForce® RTX 3080 graphics card with an image size of 512 x 512, since the employed LDM training checkpoint was optimized for this image size. Denoising was carried out by a DPM++ 2M sampler^[35]^ with a Karras scheduler^[36]^ algorithm with 25 sampling steps and sped up with Xformers^[37]^.

We generated 100 candidate images (5 batches with 20 images each) for each of our 16 scene prompts (1,600 in total), resulting in 100 image version sets. We chose 100 images to allow for enough sample variety while keeping the analysis computationally feasible, since, in a preliminary analysis, using more images resulted in qualitatively similar results. We employed negative prompts and prompt-weighting for some words to increase the likelihood of the core aspects of each scene to be generated and to reduce the likelihood of artefacts on the images (text files with our positive and negative prompts with prompt weights can be found at this OSF repository: https://osf.io/r546q/overview).

Since the core component of our pipeline is a diverse distribution of candidate images, we deemed it important to determine what level of image variability in the generated image sets is optimal for approximating the mental images. We investigated this by generating 3 different image sets (with new randomized seeds) in which we used the cfg model parameter to vary how much the diffusion model adheres to the supplied prompts. The lower the cfg parameter, the more likely it is for the model not to adhere to the prompt. This is achieved by training the model to make predictions both while conditioned on the prompt and while not conditioned on the prompt and blending between both outputs during inference (i.e., image generation) using the cfg scale parameter as a weight^[20]^. We chose cfg values 7 (medium variability), 3 (high variability) and 1 (very high variability) since they yielded visually distinct image sets with a clear image variability gradient (as demonstrated by the increasingly noisy mean images in Figure 2A). We found that, at least for our prompts, cfg values above 7 didn’t result in any significant changes in the variability of the generated image sets.

**Figure 2.**
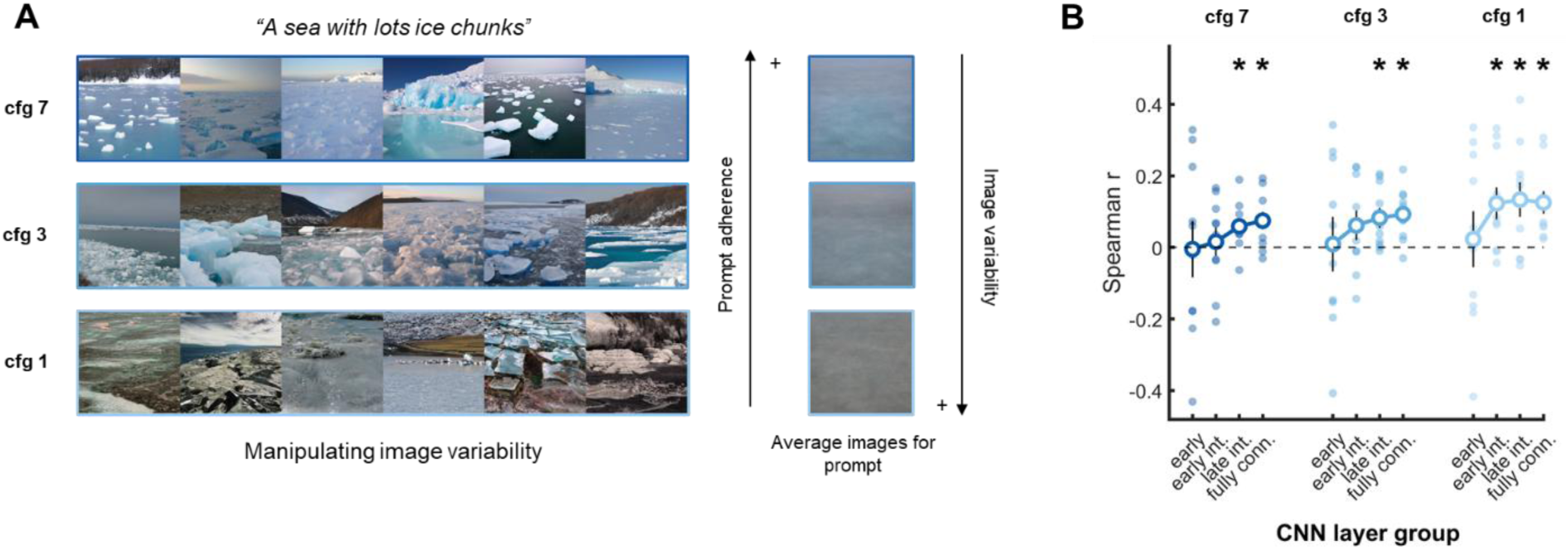
Approximating the contents of mental images. (A) A subset of exemplars from the AI-generated image sets for cfg values of 7, 3, and 1. Higher cfg values enforce greater prompt adherence while lower cfg values yield more variable images. Differences in image variability are evident from averages across 100 images obtained with each of the 3 cfg values. (B) Representational similarity between each layer group of the CNN evaluating the AI-generated candidate image sets and the participants’ alpha band representations during visual imagery. Dots are data points of individual participants. Straight lines reflect the standard error of the mean. Asterisks mark statistical significance (N = 10, right-tailed one-sample t-test, p < 0.05, FDR-corrected across layer groups).

#### Image manipulation

We investigated if we could manipulate the properties of the generated images in a way that makes them more akin to mental images. Many people report their mental images to be of a reduced sensory quality^[2,24]^, which, based on their only average vividness scores (VVIQ mean = 55, range: 48-61, compared to ^[38]^), also seems to be the case for our participants. We approximated this reduced quality, by creating degraded versions of the image set for which we found the highest correspondence with the neural imagery data (cfg 1). For each degraded image set, we applied 1 of 4 degradation types (grayscale, blurry, noisy, low contrast). We also added a control condition in which we emulated vivid mental images. We were further interested to see if we can capture some properties of mental imagery by applying a style to the images that is characterized by these properties, which we achieved by transferring the style of the cfg 1 set to 1 of 6 art styles (see Figure 3a for examples). The first set was transferred to a psychedelic style, since it envelops the images in shapeless visual flows that distort the image proportions, which might approximate what people experience during hallucinations^[21]^. The second set was transferred to a watercolor style, since we surmised its faded and blurry nature combined with its undefined outlines could capture some of the qualities of mental images, since each of these aspects had previously been described as phenomenological characteristics of mental imagery^[1]^. The third set was altered to a surrealist style which is frequently characterized by abstract, unrealistic and disproportionate scene arrangements. We wanted to determine if it can capture the more bizarre aspects of our mental imagery that have been reported^[22]^. The remaining 3 sets were then transferred to art styles that were not supposed to capture a facet of mental imagery and instead served as positive controls: 3D model, cubism, pixel art. It is worth noting that all employed styles were chosen based on intuition to some degree, since we wanted assess if our approach lends itself to such more explorative analyses of the properties of mental images.

**Figure 3.**
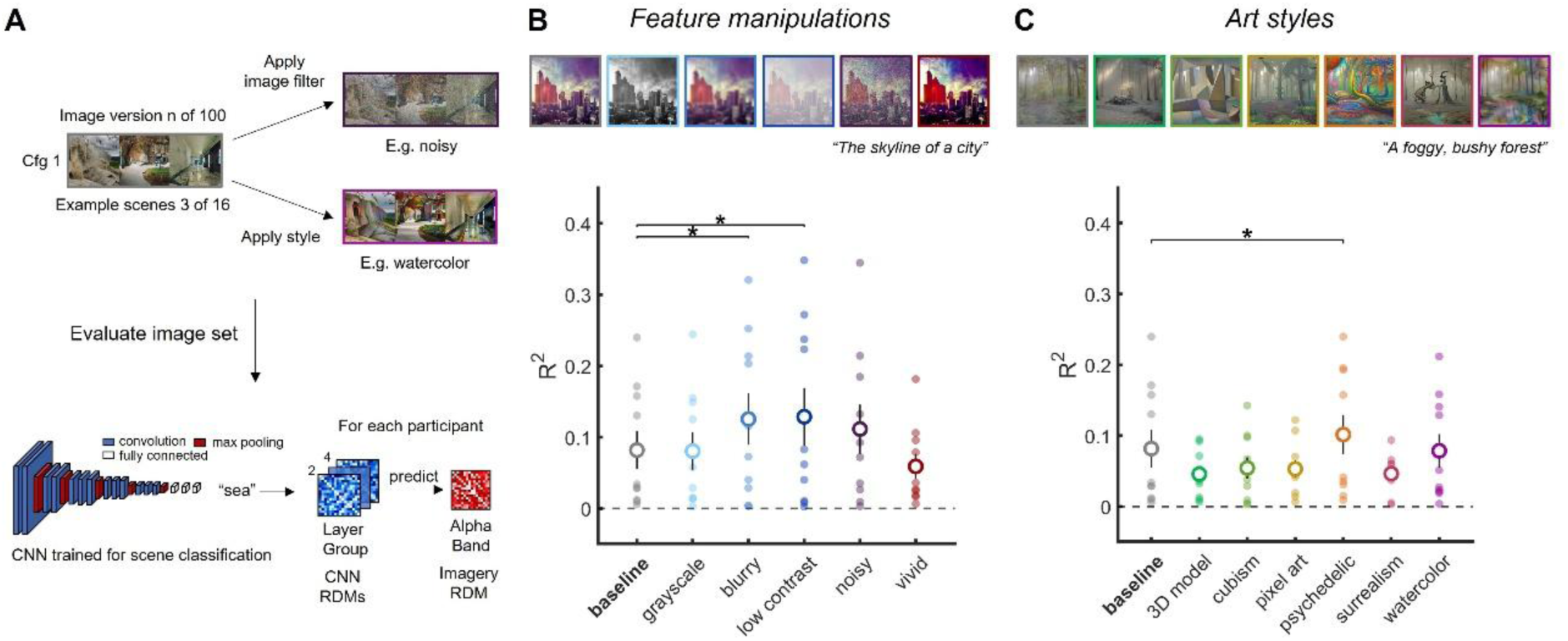
Approximating the qualities of mental images. (A) Property manipulation pipeline. We manipulated the properties of the cfg 1 image set, for which we found the highest representational alignment between the evaluator CNN and our participants’ imagery-related neural representations, to examine if targeted image alterations further enhance this representational alignment. In the image feature manipulation, we either applied 1 of 4 types of image degradation (grayscale, blurry, low contrast, noisy) to emulate the reduced sensory quality of mental images or, as a control, made the images more vivid (high contrast, high saturation). In the style manipulation, we employed a LDM fine-tuned for style transfer to transfer the style of the image set to 1 of 6 art styles (3D model, cubism, pixel art, psychedelic, surrealism, watercolor). For each image set (the baseline set and all manipulated sets), we again extracted RDMs from the evaluator CNN. We then used RDMs from layer groups 2-4 (early intermediate to fully connected), for which we previously found representational alignment with participants’ alpha activity, to predict the alpha band RDM of each participant with a regression model. For each fit regression, we computed the explained variance (R²), averaged it across participants and compared it to the explained variance of the baseline image set (cfg 1). (B) Representational alignment between evaluator CNN and participants’ alpha band representations for the image feature manipulation with a curated visualization of the image manipulations for 1 of the 1,600 images in the baseline set. Degrading the images by adding Gaussian blur or reducing the contrast significantly improved representational alignment compared to baseline. (C) Representational alignment between evaluator CNN and participants’ alpha band representations for the style manipulation with a curated visualization of the image manipulations for 1 of the 1,600 images in the baseline set. Transferring the images to a psychedelic style which envelops the images in shapeless visual flows that distort the image proportions resulted in significantly enhanced representational alignment compared to baseline. Dots are data points of individual participants. Straight lines reflect the standard error of the mean. Asterisks mark statistical significance (N = 10, right-tailed paired Wilcoxon signed-rank test, p < 0.05, FDR-corrected across comparisons).

We manipulated the image features with in-built MATLAB image processing functions. The blurred image set was smoothed with a 6 mm full-width at half maximum (FWHM) gaussian kernel. For the noisy image set, gaussian noise (mean = 0, variance = 0.3) was added. The grayscale image set was created by reducing the color space from RGB to grayscale. The low contrast images were created by limiting the intensity range of the images between 0.6 and 0.9. We emulated vivid mental images by increasing the contrast by mapping the intensity range from 0.2-0.9 to 0-1 via histogram stretching and multiplying the saturation by a factor of 1.5 and clipping the values at 1 to not exceed the saturation range.

The style manipulation was conducted using DreamShaper 8 (https://civitai.com/models/4384/dreamshaper) as the base model (cfg scale = 7, denoising strength = 0.6), a SD-1.5-based architecture fine-tuned for art style transfers. For each image in the set, we first had ControlNet (v1.1.455; ^[39]^), a neural network that allows for additional spatial conditioning of the LDM, apply a Canny edge detection preprocessor to create an edge map of each image (using default parameters). This allowed us to roughly maintain the structure of the original image by conditioning the output image on the edge map with a high weight. The art style was then altered with the SDXL Style selector extension (https://github.com/ahgsql/StyleSelectorXL) for Automatic1111. This extension can change the art style of a reference image by adding positive and negative prompts for 300 selectable art style presets. We chose the following presets for the respective art styles: Psychedelic, Watercolor, Surrealist, 3D Model, Cubist and Pixel Art. Finally, the output image was conditioned on the original image, the style-specific positive and negative prompts as well as the ControlNet edge map. The sampler and scheduler were identical to those in the previous image generation pipeline, but we reduced the number of sampling steps to 20 to speed up processing. For a visualization of all 14 image sets we used for analysis, see supplementary Figures S4-S18. The full image sets can be found on Hugging Face: https://huggingface.co/datasets/RicoStecher/Imagery_Approximation_Image_Sets.

#### CNN pipeline

Next, we assessed how good of an estimate the generated image sets are by using a CNN to computationally simulate visual cortex responses to the images and comparing them to actual visual content representations in the alpha rhythms during imagery via RSA^[16]^.

The representations of CNNs trained for image recognition offer state-of-the-art predictions of visual cortex representations during perception^[40,41]^. Xie et al.^[5]^ showed that, if supplied with images of the imagined visual contents, they can also be used to model reactivated visual content representations encoded in the alpha rhythms of participants forming a mental image. Thus, if our generated image sets are sufficient as estimates of the imagined visual contents, there should be a correspondence between the visual content representations in the alpha rhythms during imagery and the visual CNN representations when classifying the candidate images.

The analysis was conducted using VGG-16^[17]^ trained for scene recognition on the Places365 image set^[18]^. We employed this model, since it is one of the best models of the visual feedforward cascade among CNNs^[42]^ and since Xie and colleagues^[5]^ also used a VGG architecture to successfully model rhythmic content representations during object imagery.

The images were downsampled to 224 x 224 pixels to match the VGG-16 input layer size. For each of the 100 image sets, we evaluated the 16 images within each image set with the VGG-16 model, obtained the unit activations and constructed RDMs at each layer. This was achieved by computing correlation distances (1 - Pearson correlation) between the layer unit activations patterns elicited by each pair of generated scene images and assembling them into a 16 x 16 matrix. We then averaged the RDMs at each layer across all 100 image sets to get an average estimate of the discriminability among generated images across the entire distribution. Since there is a high autocorrelation between RDMs of neighboring CNN layers, we divided the 16 convolutional and fully connected layers into 4 layer groups and averaged the RDMs in each layer group in order to improve power^[5]^. The early layer group consisted of the first 4 convolutional layers, the early intermediate layer group of spanned convolutional layers 5-10 and the late intermediate layer group convolutional layers 11-13. The fully connected layer group consisted of all 3 fully connected layers (layers 14-16). We repeated this approach for all 3 cfg values.

When conducting the image manipulation analysis, we used the same pipeline to create layer group RDMs for each image set in the image feature manipulation and the style manipulation (see Figure 3A).

#### EEG analysis pipeline

In order to access the rhythmic neural representations of the imagined scene contents to be modelled by the CNN, we created RDMs at every EEG frequency from 8-13 Hz (spanning the entire alpha band). Initial decoding analyses revealed robust stimulus information in the alpha band, in line with previous findings (^[5,7]^; see supplementary Figure S2). All multivariate analyses on the neural data were conducted in CoSMoMVPA^[43]^ in MATLAB 2022a.

Due to the non-stationary nature of the EEG signal, EEG response pattern distributions can vary across sessions to such a degree that it can limit pattern-based analyses^[44]^.

In order to remedy this, we employed a domain adaptation approach when creating the RDMs. Domain adaptation is a common machine learning practice in which differing distributions in multiple datasets (or *“domains”*) are aligned through various algorithms^[45]^. Here, we used semi-supervised maximum independence domain adaptation (sMIDA; ^[19]^) to map the original power pattern space across all electrodes at each frequency to a feature space that is maximally independent from all sessions to deal with session-by-session variation and maximizes the discriminability among scenes. We modelled each of the 10 sessions as a discrete domain. Since this approach uses label information to maximize discriminability, we estimated the feature space anew for each of the 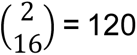 pairs of imagined scenes to maximize the neural discriminability for each pair. We utilized the sMIDA algorithm implemented in the MATLAB domain adaptation toolbox (https://de.mathworks.com/matlabcentral/fileexchange/56704-a-domain-adaptation-toolbox?requestedDomain=). We used the following hyperparameters: linear kernel, m = 8, gamma = 0.5, sigma = 10, mu = 0.01. RDMs were then created by averaging all 270 trials of each stimulus across all 10 sessions to increase power, calculating a correlation distance between the average activation patterns in the domain-independent feature space elicited by each imagined scene within each scene pair and assembling the distances into 16 x 16 RDMs. We further boosted the SNR by identifying at which frequency in the alpha band the imagined scenes could be discriminated the best for each participant and using the RDM at that frequency for analysis. We achieved this by averaging the distances in the bottom triangle of each RDM for each frequency as a measure of general neural discriminability among the imagined scenes and pinpointing the frequency in the alpha band for which this was maximal.

#### RSA modelling

Finally, we assessed the representational similarity between the alpha band RDM of each participant during scene imagery and the CNN RDM of each layer group by calculating a Spearman correlation. Correlations at each layer group were averaged across participants. We repeated this with the image sets of each cfg value. If there is a positive average correlation with a particular layer group, that would suggest that: 1.) The generated image set was sufficient as an approximation of the participants’ mental images and activated visual representations in the CNN that are similar enough to the actual representations of the imagined visual contents encoded in the neural rhythms of the participants to cause a correspondence. 2.) Since the CNN extracts visual features from an image in a hierarchically convergent manner (i.e., the deeper the layer, the more complex the represented visual features), the depth of the layer group at which we find significant correlations provides evidence for the level of complexity of the visual features of the visual contents encoded in the rhythms.

To investigate if the properties of the generated images can be manipulated to make them more alike the participants’ actual mental images, we fit a regression model for every image-feature-manipulated image set and for every style-transferred image set, using the layer group RDMs for which we originally found a correspondence with the neural imagery data (early intermediate, late intermediate and fully connected) as predictors for the alpha band RDM of each participant. We calculated the explained variance (R²) for each participant, averaged it across participants and compared it to the average R² of the unaltered cfg 1 image set, to see if it improves the fit with the neural imagery data (see Figure 3A & C). It is worth noting that the resulting R² values in this approach cannot be interpretated against 0, but comparing the R² for different manipulations yields an estimate of the fit improving or reducing when a manipulation is performed.

## Statistical inference

For the initial assessment of the approximations the participants’ mental images, we used a right-tailed one-sample t-test (N = 10, p < 0.05) to assess if the correlation between the imagery alpha band RDM and each CNN layer group RDM is greater than 0. The uncorrected p-values were FDR-corrected across CNN layer groups.

For the image manipulation analysis, we used a right-tailed paired Wilcoxon signed rank test to compare the R² of every image manipulation type (i.e., image feature manipulation or art style) to the R² of the baseline image and FDR-corrected the p-values across all such comparisons separately for the image feature manipulation and style manipulation.

## Code and Data Availability

Analysis code can be accessed from this GitHub repository: https://github.com/DKaiserLab/imagery_approximation

Materials and data can be accessed from this OSF repository: https://osf.io/dxnhm/overview

## Additional information

The authors declare no conflicts of interest.

## Author contributions

**R.S.:** Conceptualization, Methodology, Data curation, Investigation, Formal analysis, Visualization, Writing — original draft, Project administration, Writing — review and editing

**D.K.:** Conceptualization, Methodology, Supervision, Project administration, Funding acquisition, Writing — review and editing

## Acknowledgements

We would like to thank Tanja John for organizing and conducting the data collection of most of this very extensive EEG dataset, but also extend our thanks to Max Bardelang, Malaika Alphonsus, Marius Geiss, Tugce Dalmis, und Pietra Pacheco Alves for helping with data collection. R.S. and D.K. are funded by an ERC Starting Grant (PEP, ERC- 2022-STG 101076057). This work is further supported by the DFG under Germany’s Excellence Strategy (EXC 3066/1 “The Adaptive Mind”, project number 533717223). Views and opinions expressed are those of the authors only and do not necessarily reflect those of the funders. Neither the funders nor the granting authority can be held responsible for them.

## Notes

https://github.com/DKaiserLab/imagery_approximation

https://osf.io/r546q/overview

https://huggingface.co/datasets/RicoStecher/Imagery_Approximation_Image_Sets

### Competing Interest Statement

The authors have declared no competing interest.

### Summary of Updates

Fixed panel letters in Figure 3.

