## Supplement for "A neuro-computational approximation of the qualities of mental images"

**Supplementary Information**

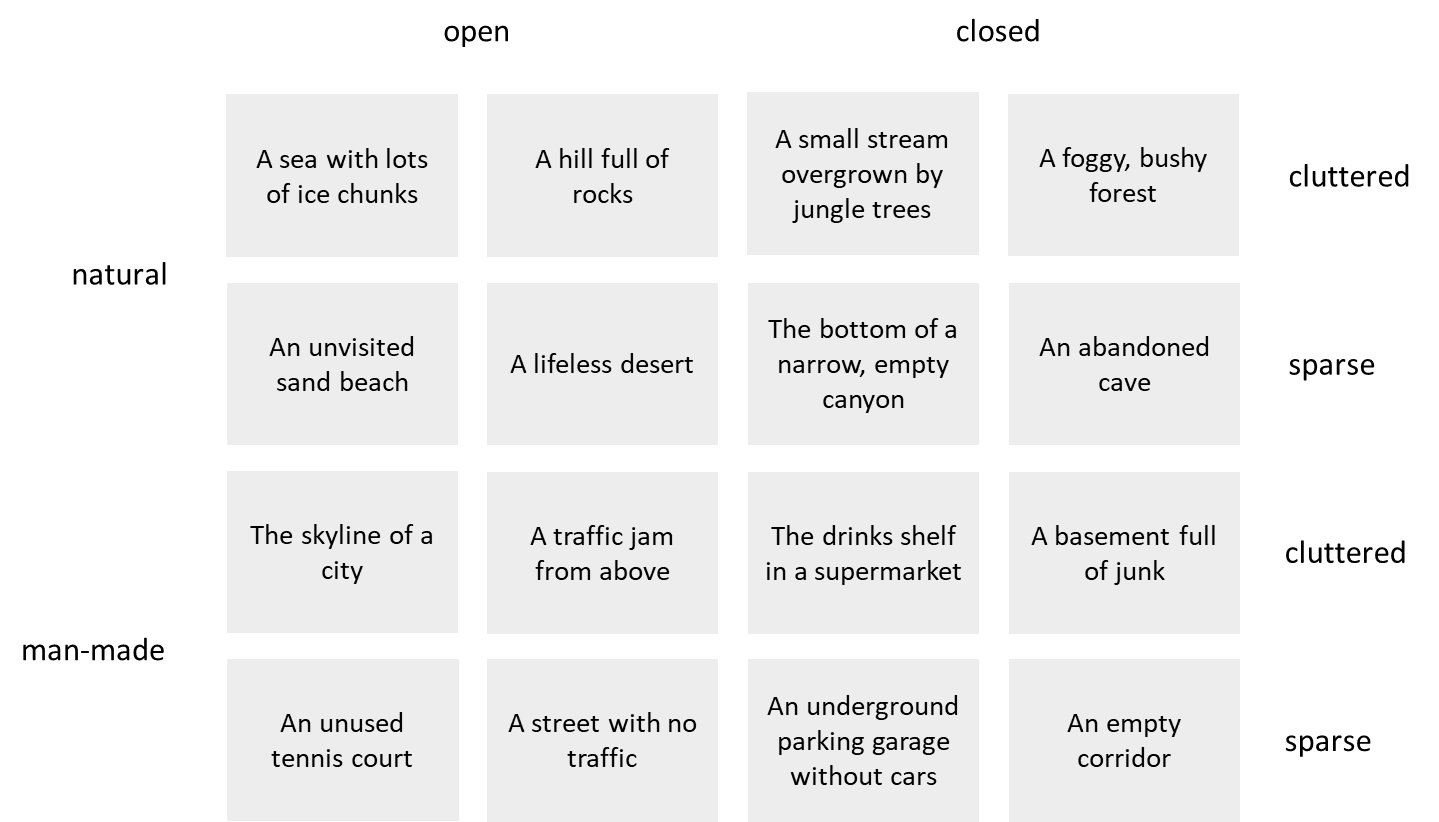

**Figure S1. Scene prompts with scene property categories.**

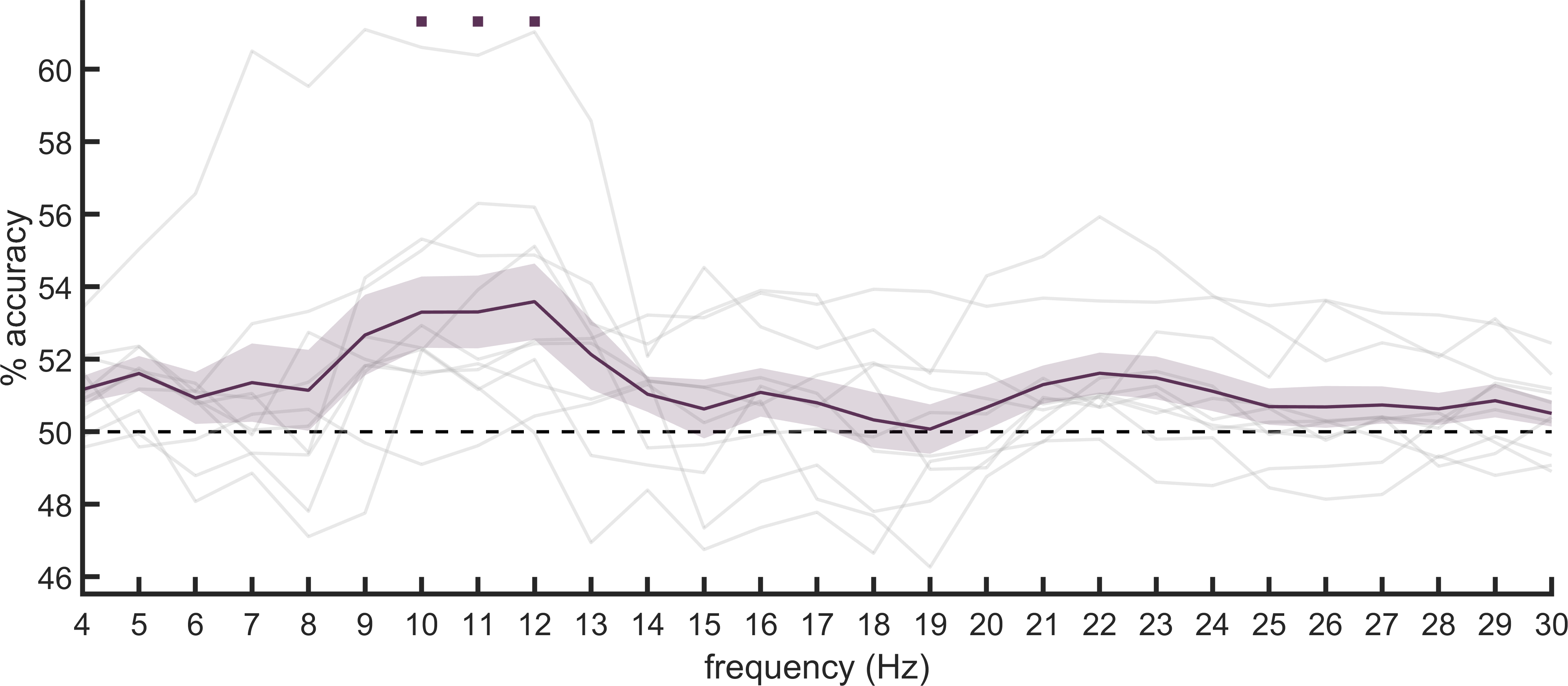

**Figure S2. Neural discriminability of imagined scenes across frequencies.** To assess if the imagined scenes could be discriminated alpha power patterns, we trained LDA classifiers to discriminate between every pair of imagined scenes based on the domain-adapted scalp power patterns at each frequency from 4-30 Hz (adding spanning the theta and beta bands for reference) and then averaged across scene pairs. The sMIDA space was estimated in the training set and applied to the test set for each of the 9 cross-validation folds. We then created 11 pseudotrials for each stimulus in each of the folds by randomly selecting a trial of that stimulus for each of the 10 sessions (ensuring that the trial pools were kept independent across folds) and averaging them. The classifiers were then trained on the pseudotrials in a leave-one-fold-out scheme. We repeated this for 100 permutations and averaged the accuracies across permutations. As expected, the imagined scenes could be discriminated from alpha band activity. Gray lines indicate decoding accuracies of individual participants. Squares indicate significant above-chance decoding (Threshold-Free Cluster Enhancement, 10,000 sign permutations, FWER-corrected across frequencies). Shaded areas mark the standard error of the mean.

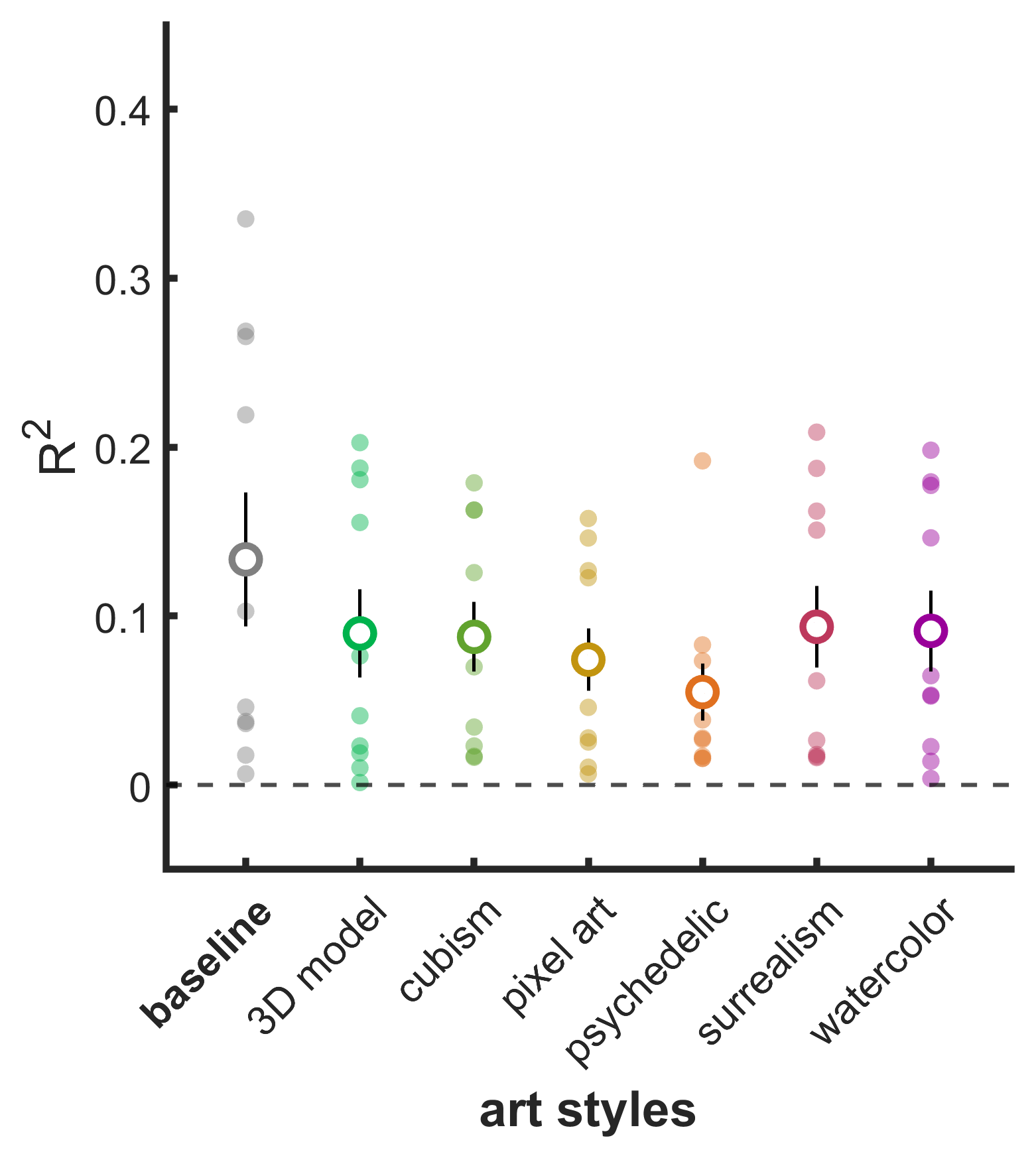

**Figure S3. CNN-alpha-band correspondence with a CNN trained on stylized object images.** It is possible that the style manipulation analysis was limited by the stylized images being out-of-distribution for the CNN we employed, since it was trained on realistic scene images. For that reason, we investigated if utilizing a model that was trained on stylized images would improve the CNN-brain correspondence for the style-transferred image sets. We utilized VGG-16 trained on Stylized-ImageNet (Geirhos et al., 2019). However, this yielded no such improvements and the model did not replicate the previously observed advantage for images in a psychedelic style (see Fig. 3).

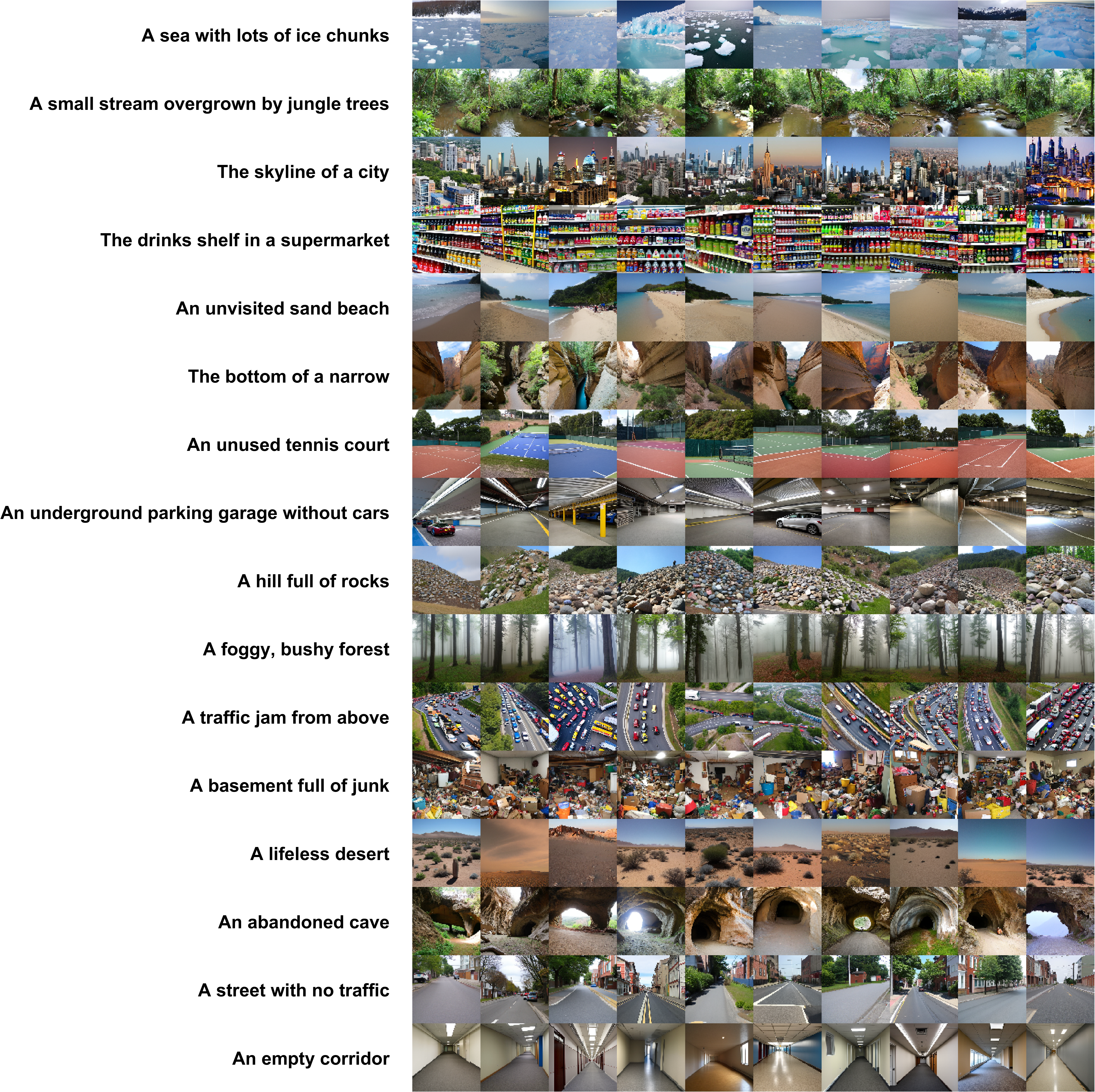

**Figure S4. Example stimuli (first 10 versions of the 16 scenes) for the cfg 7 image set.**

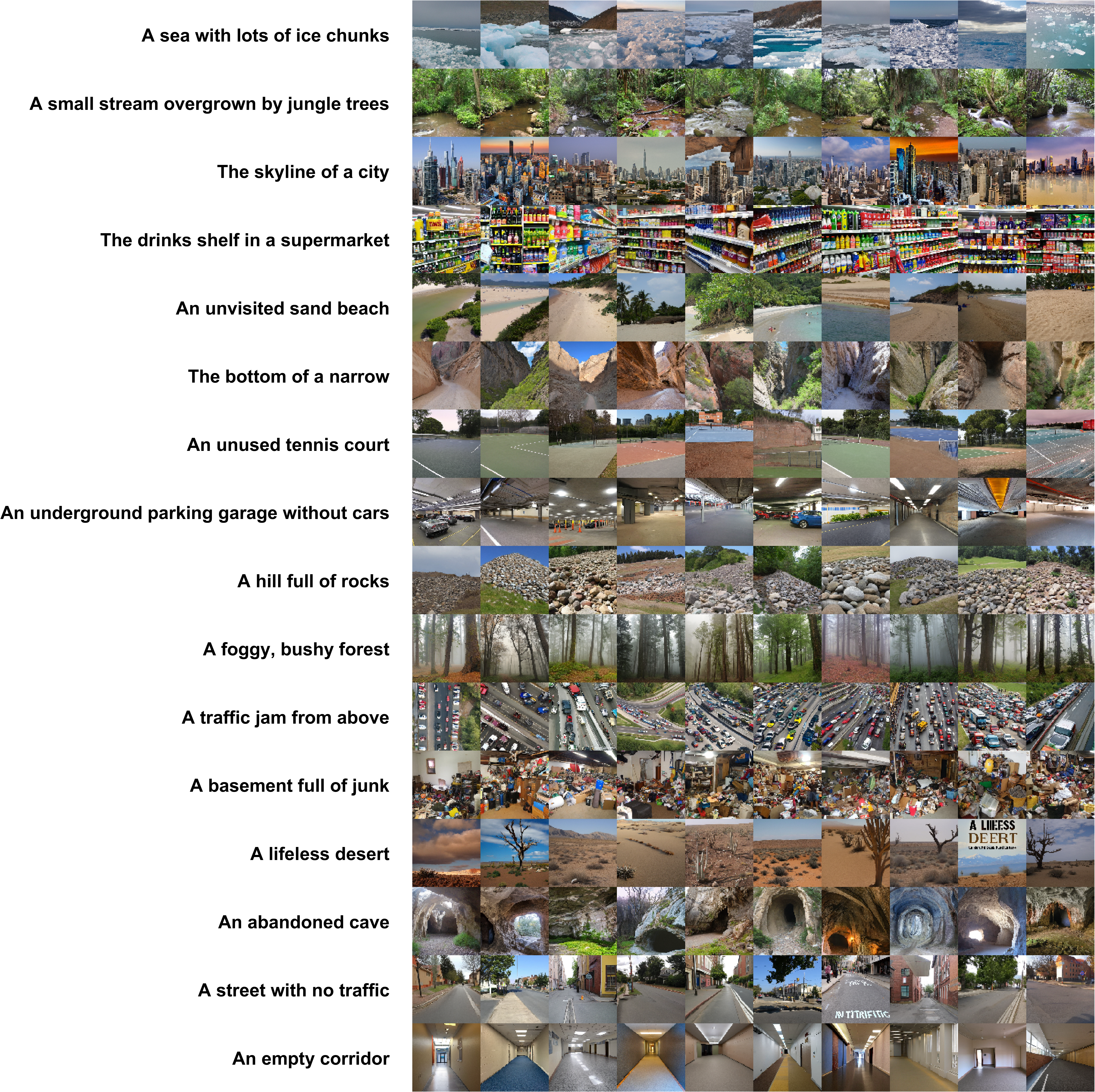

**Figure S5. Example stimuli (first 10 versions of the 16 scenes) for the cfg 3 image set.**

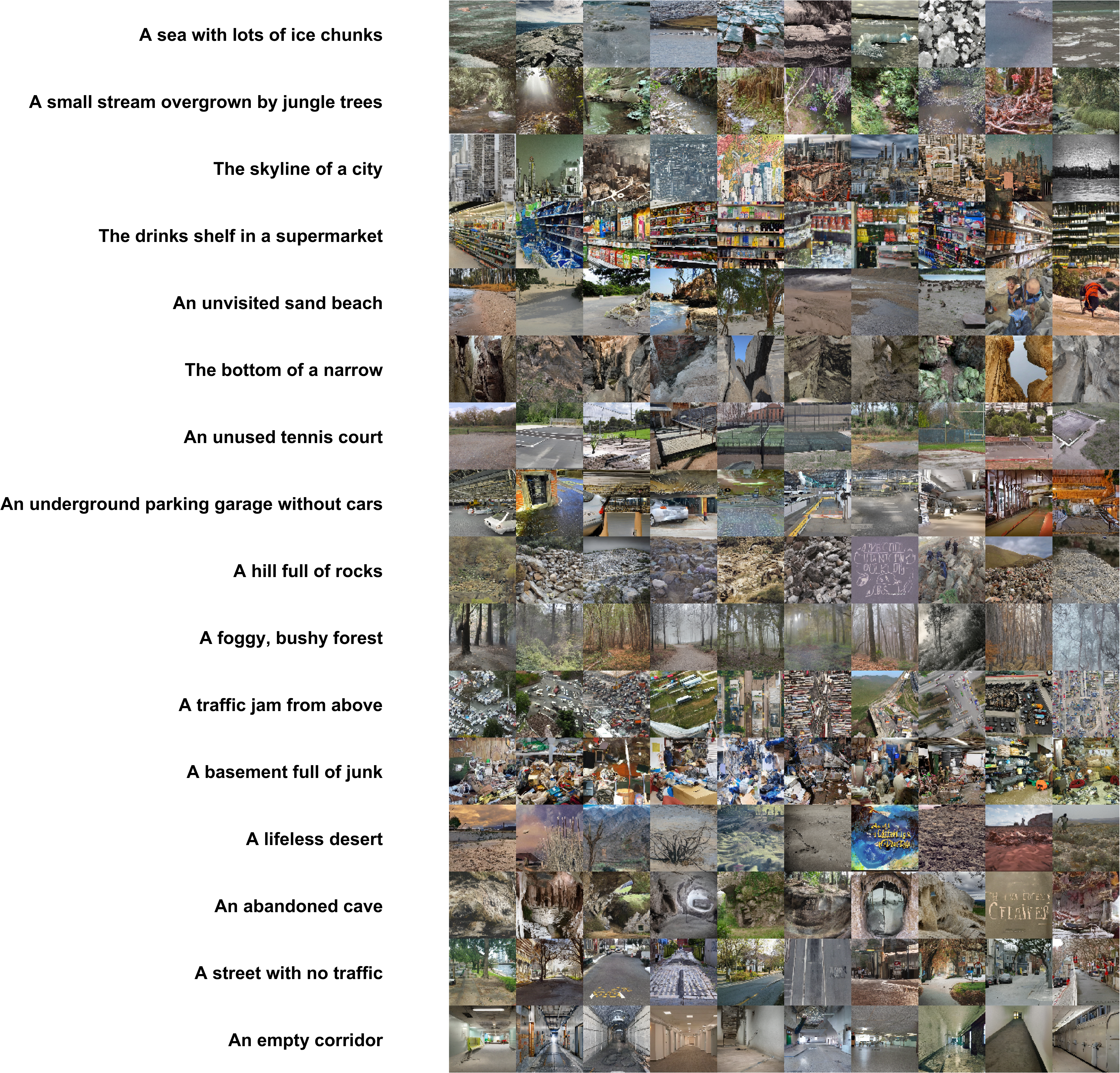

**Figure S6. Example stimuli (first 10 versions of the 16 scenes) for the cfg 1 image set (baseline set for the image manipulation analysis).**

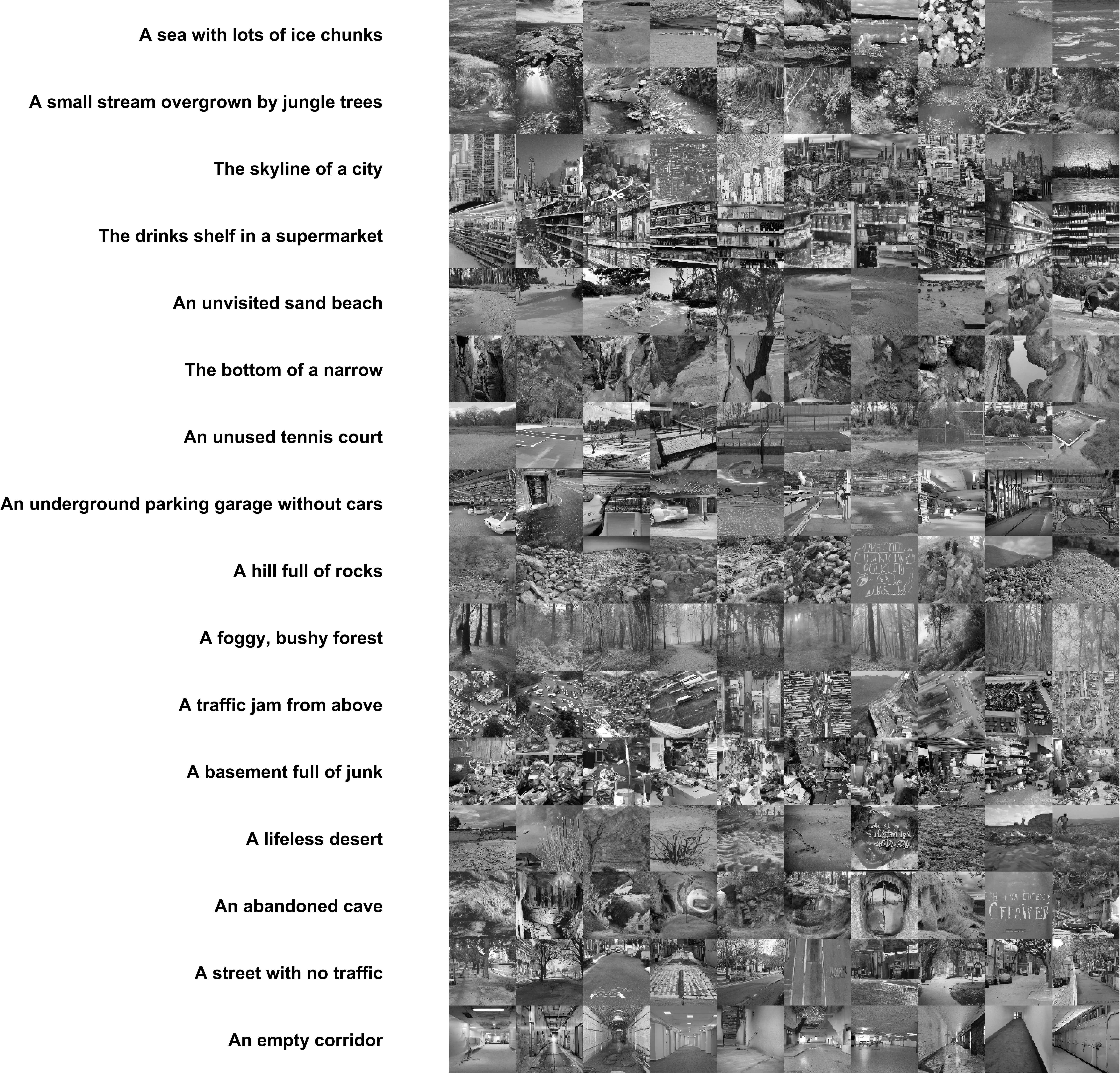

**Figure S7. Example stimuli (first 10 versions of the 16 scenes) for the grayscale manipulation.**

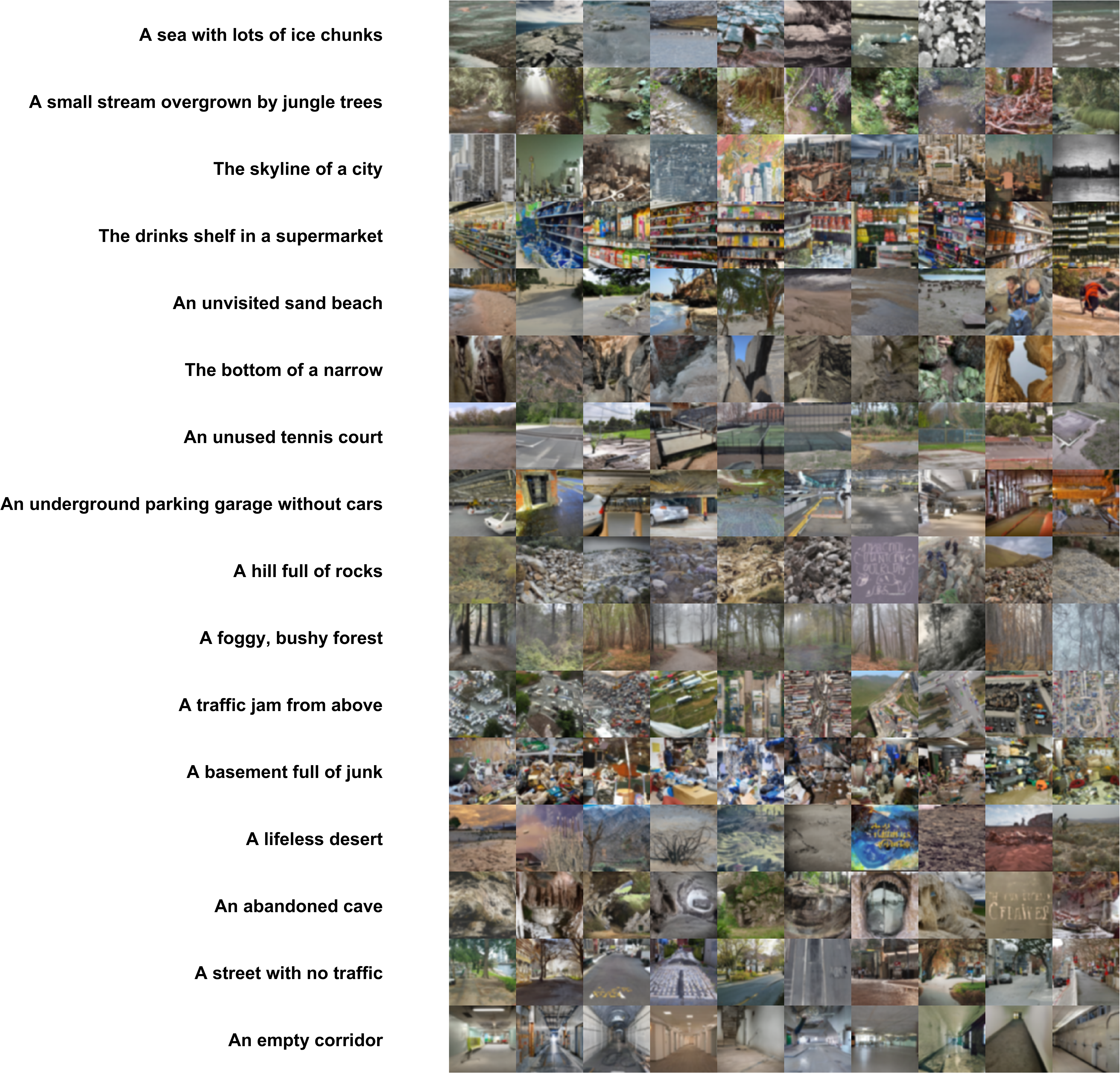

**Figure S8. Example stimuli (first 10 versions of the 16 scenes) for the blurry manipulation.**

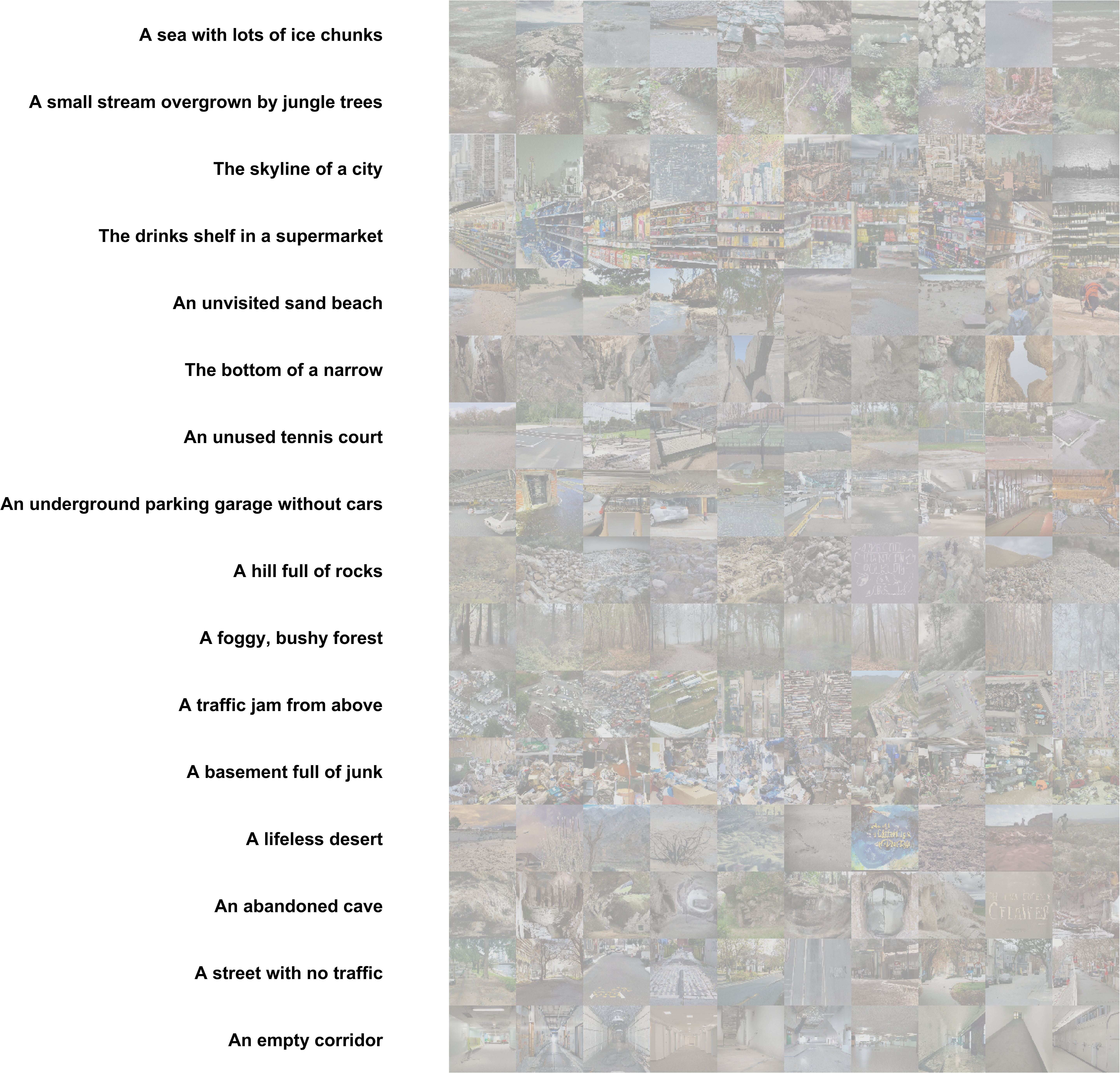

**6. Example stimuli (first 10 versions of the 16 scenes) for the low contrastmanipulation.**

**Figure S9. Example stimuli (first 10 versions of the 16 scenes) for the low contrast manipulation.**

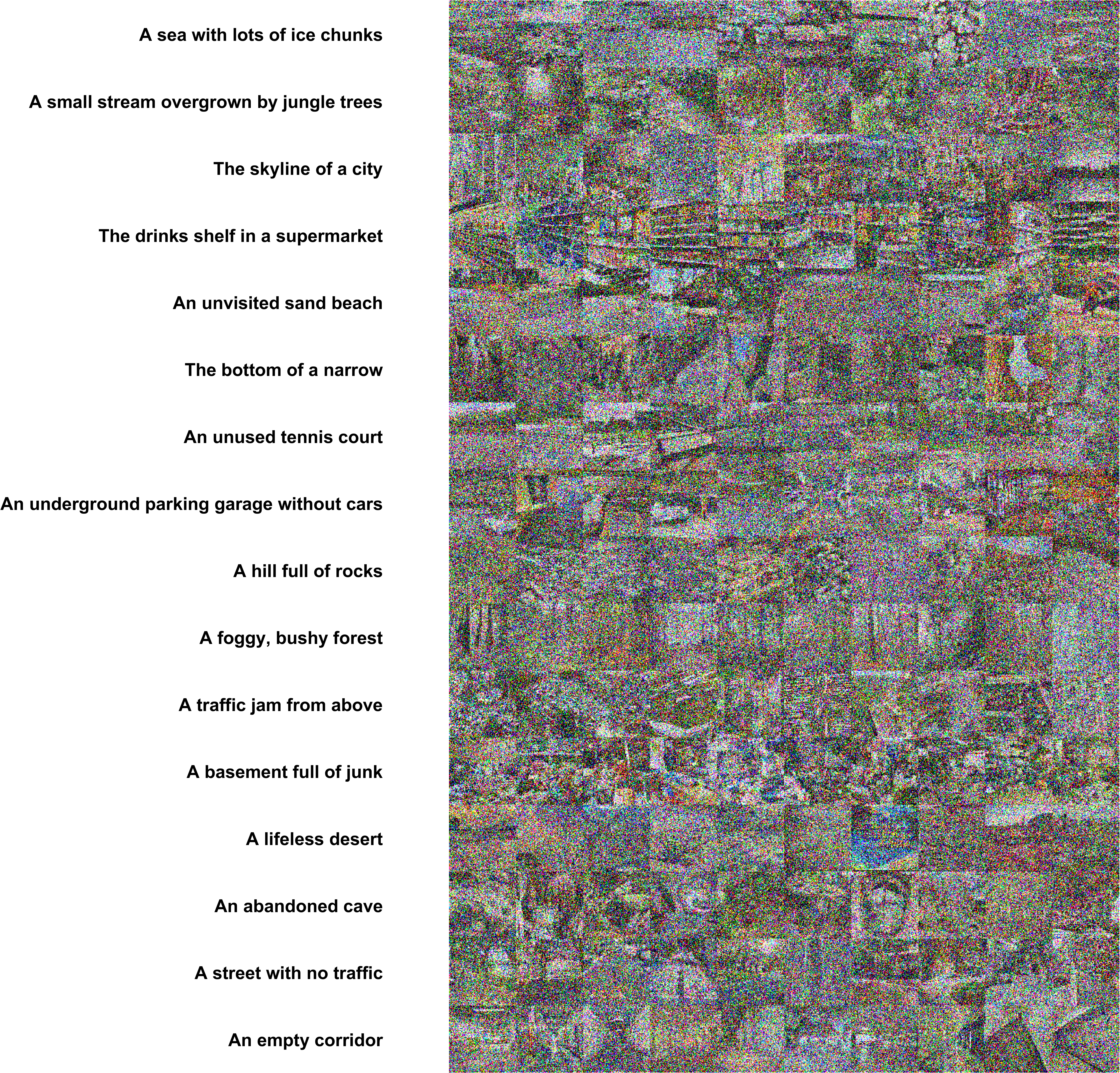

**Figure S10. Example stimuli (first 10 versions of the 16 scenes) for the noisy manipulation.**

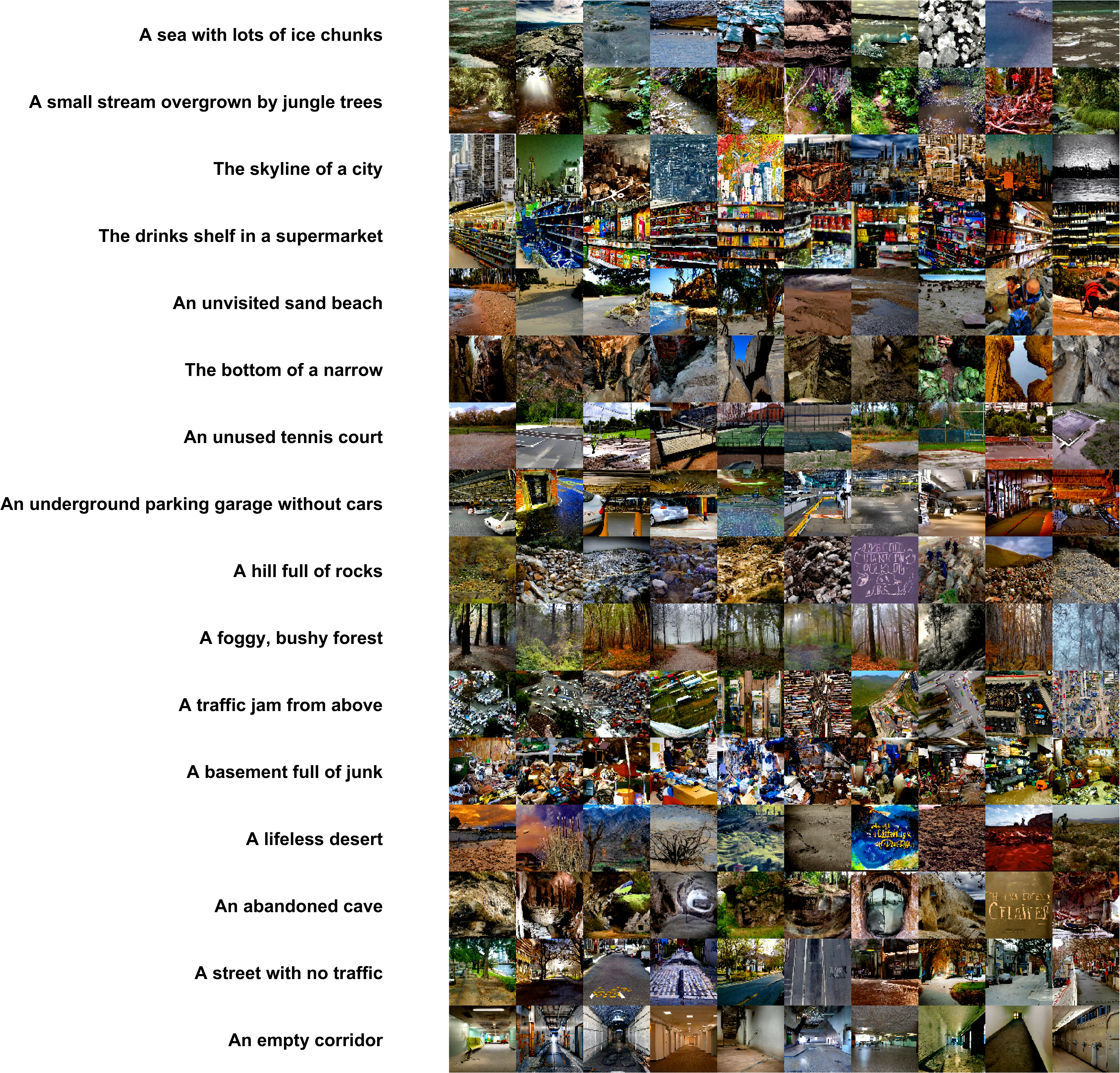

**Figure S11. Example stimuli (first 10 versions of the 16 scenes) for the vivid manipulation.**

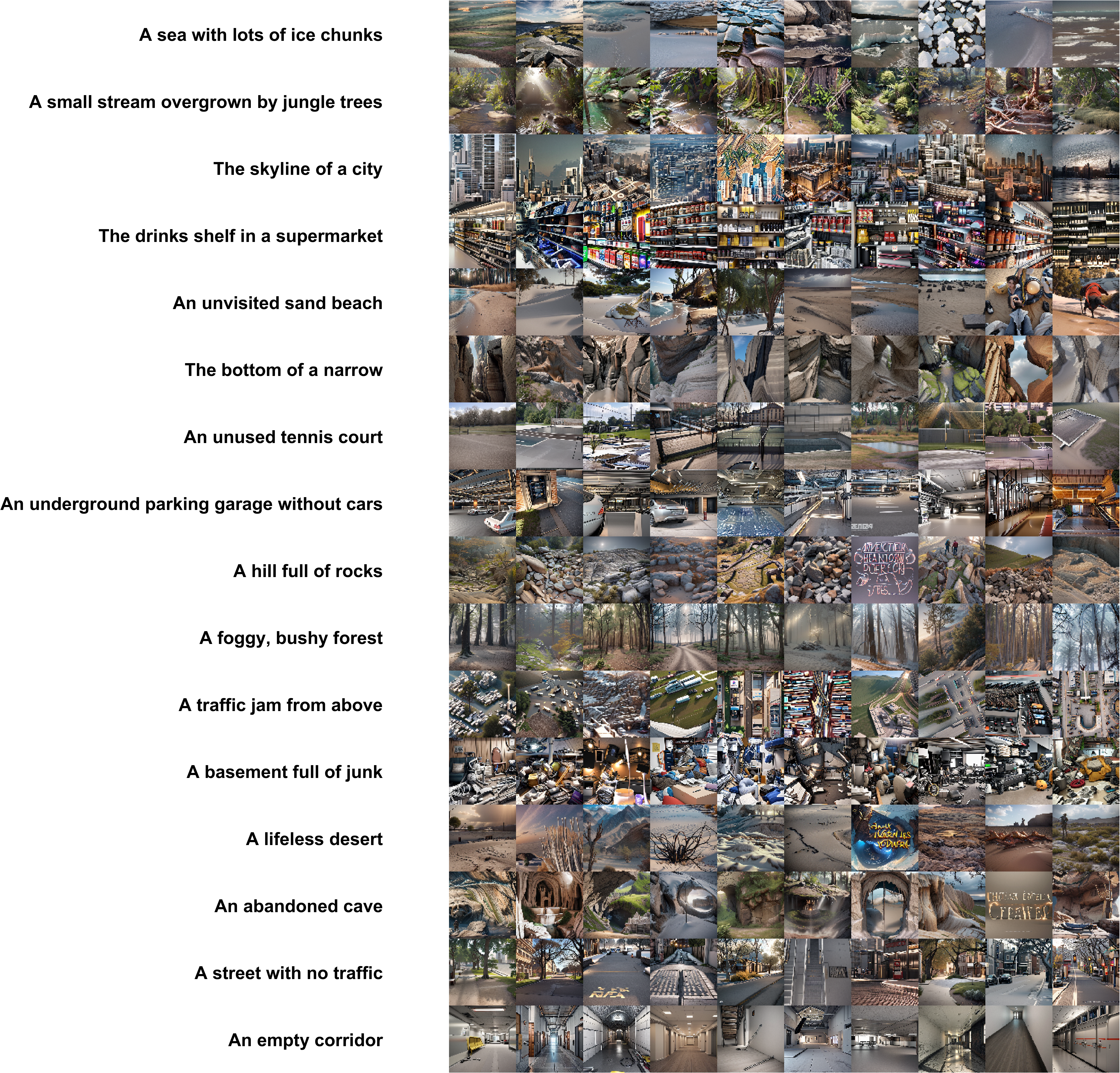

**Figure S13. Example stimuli (first 10 versions of the 16 scenes) for the 3D model manipulation.**

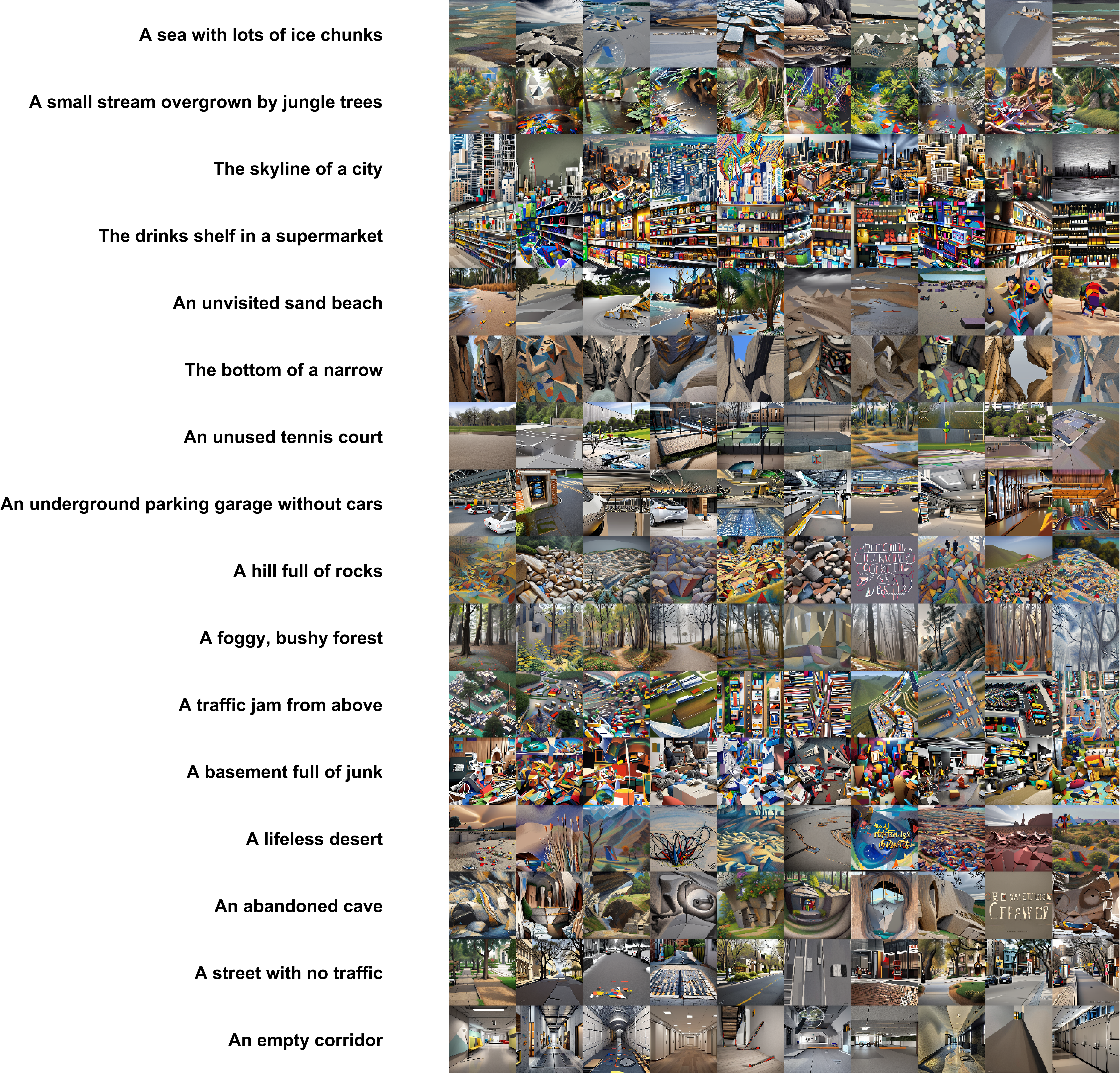

**Figure S14. Example stimuli (first 10 versions of the 16 scenes) for the cubism model manipulation.**

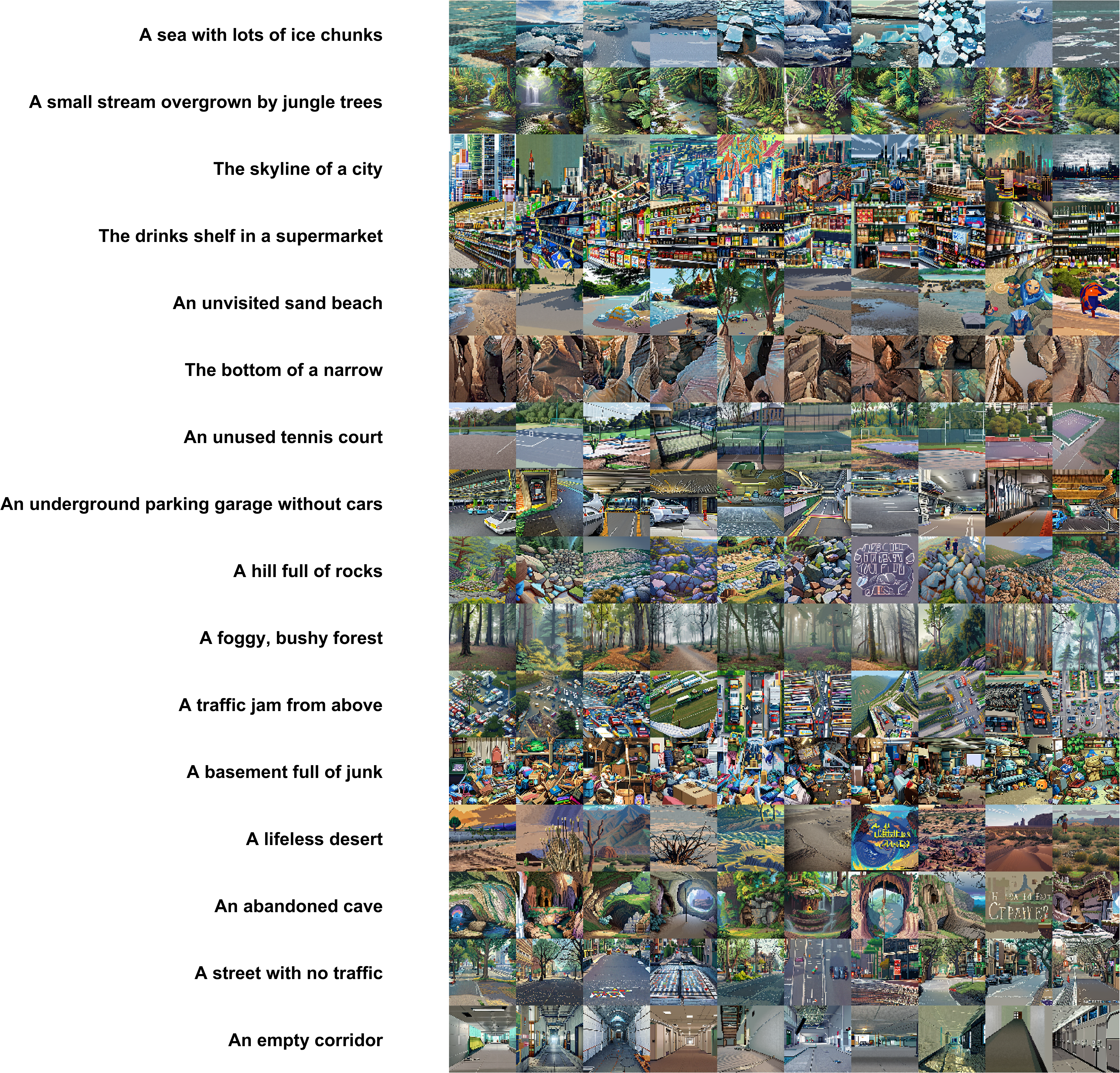

**Figure S15. Example stimuli (first 10 versions of the 16 scenes) for the pixel art manipulation.**

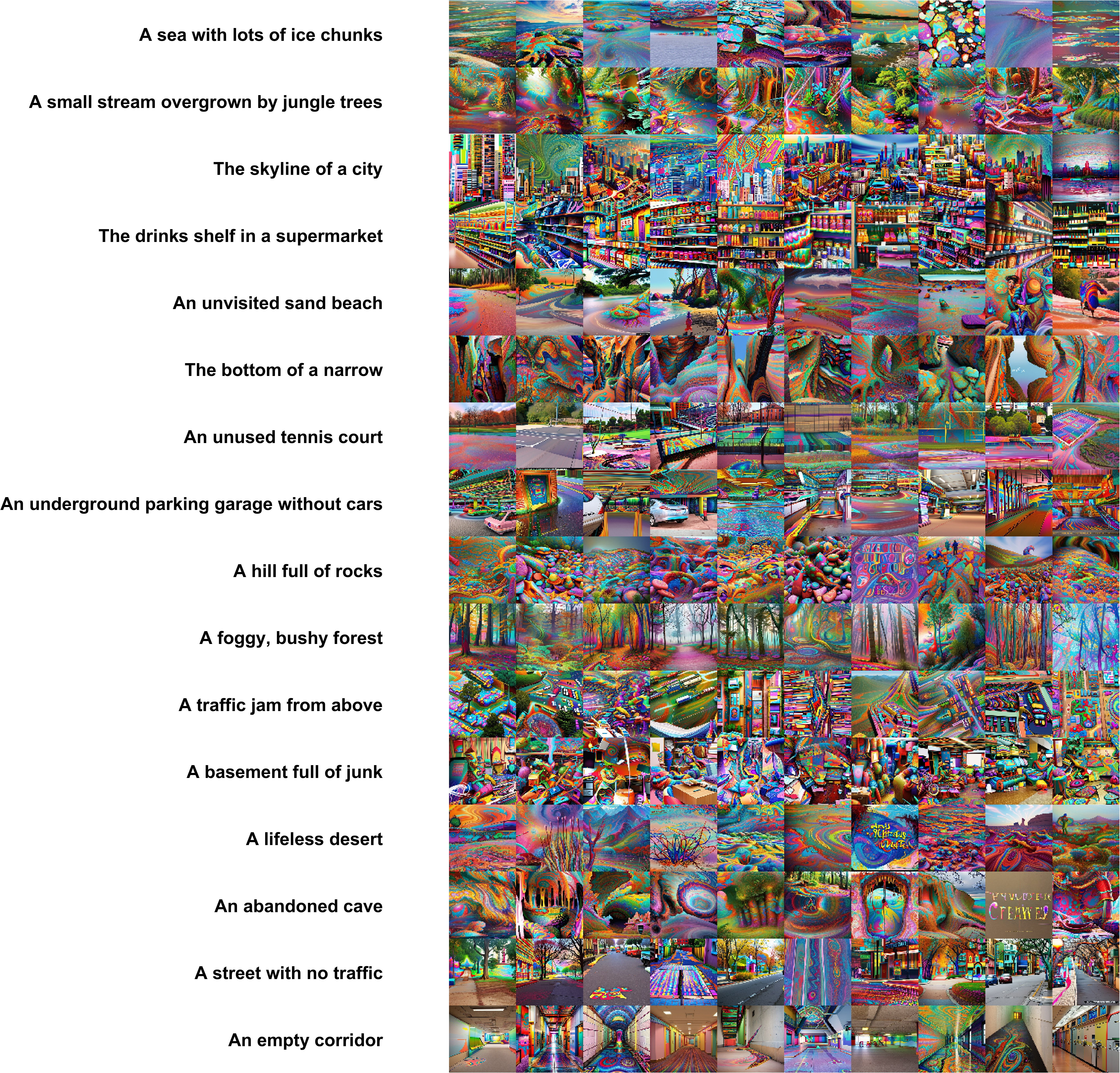

**Figure S16. Example stimuli (first 10 versions of the 16 scenes) for the psychedelic model manipulation.**

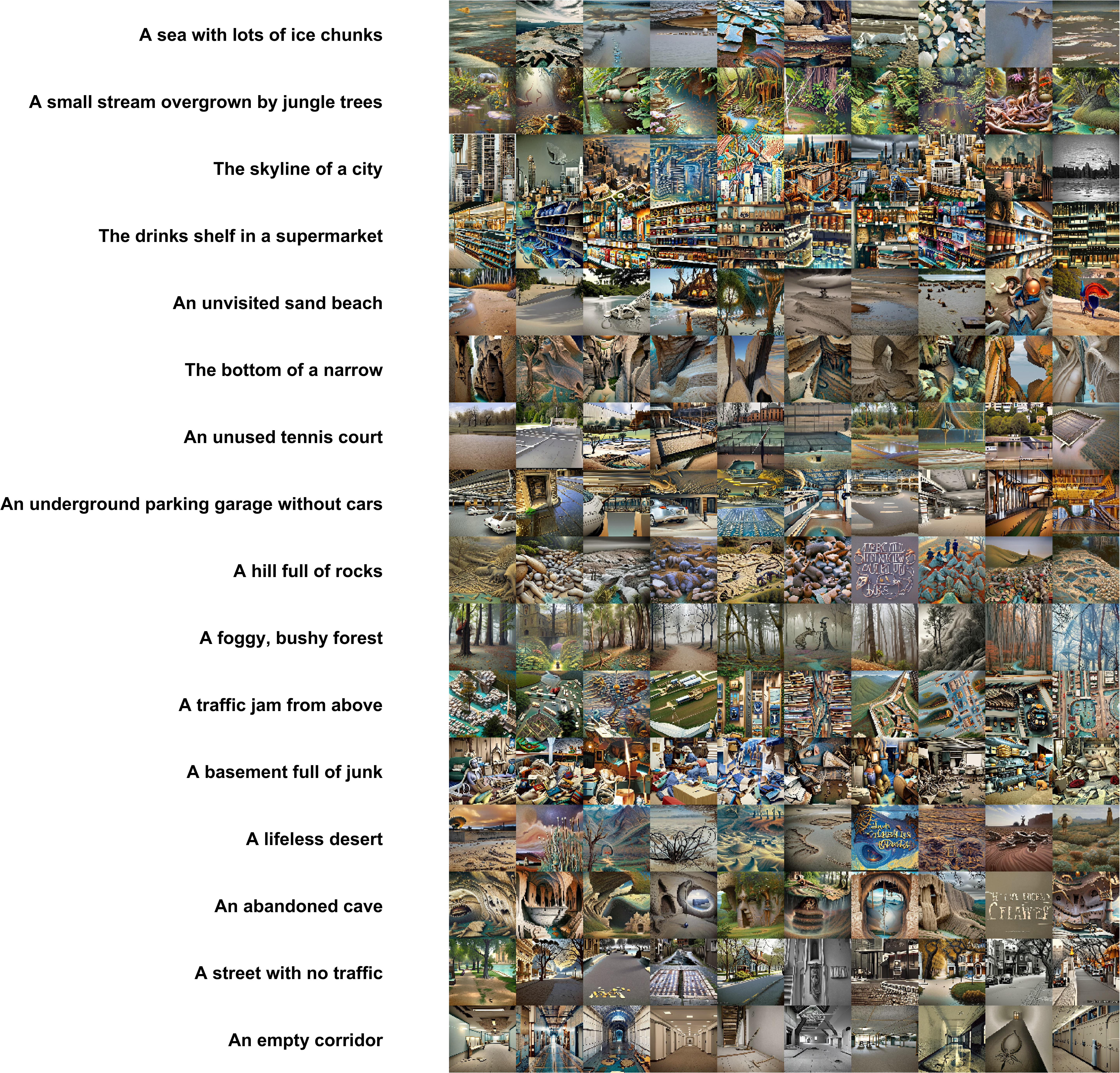

**Figure S17. Example stimuli (first 10 versions of the 16 scenes) for the surrealism manipulation.**

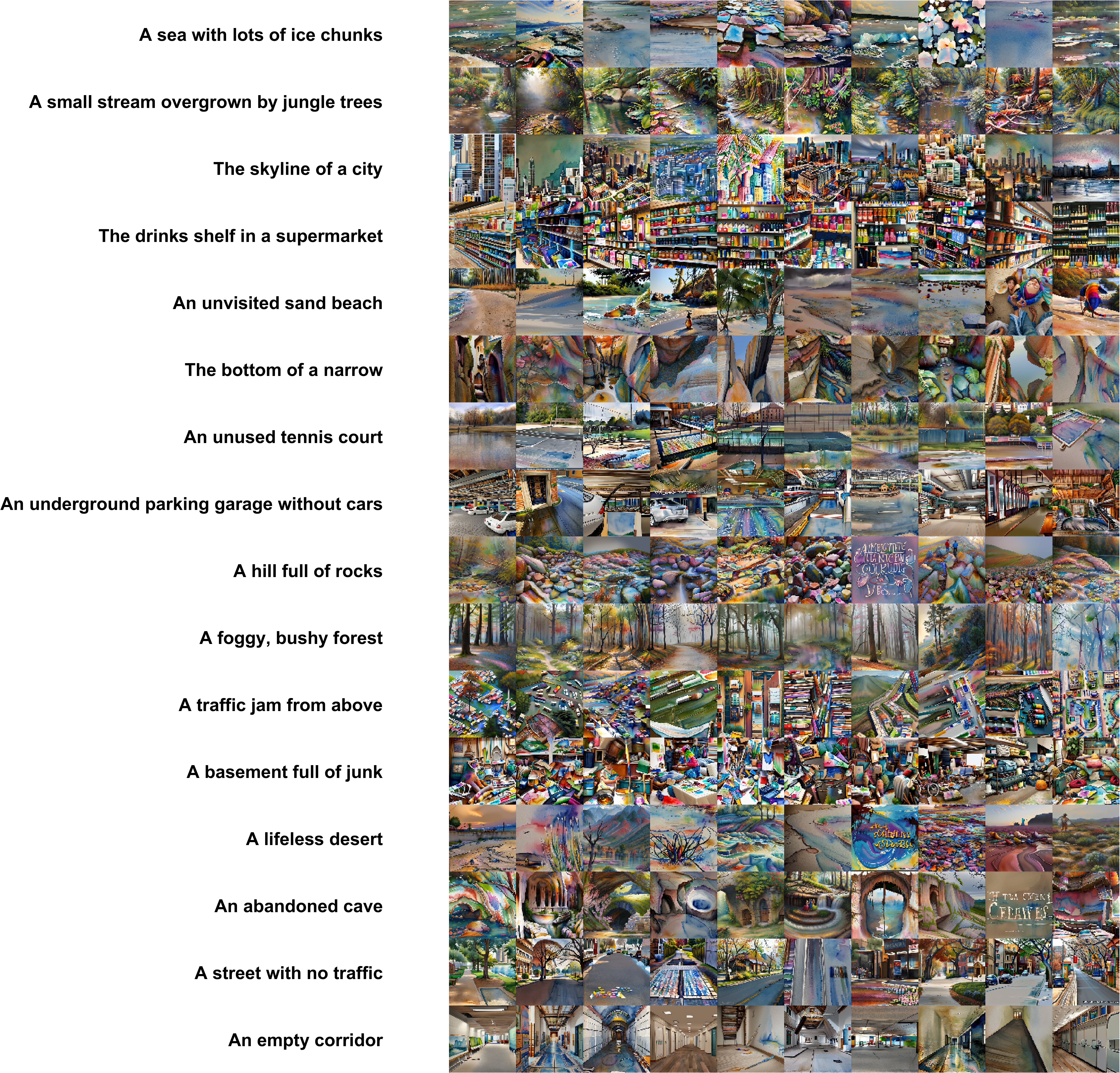

**Figure S18. Example stimuli (first 10 versions of the 16 scenes) for the watercolor manipulation.**
